# Letter: Cancer-independent, second somatic *NF1* mutation of normal tissues in neurofibromatosis type 1

**DOI:** 10.1101/2024.09.09.611411

**Authors:** Thomas R. W. Oliver, Andrew R. J. Lawson, Henry Lee-Six, Anna Tollit, Hyunchul Jung, Yvette Hooks, Rashesh Sanghvi, Matthew D. Young, Timothy M. Butler, Pantelis Nicola, Taryn D. Treger, G. A. Amos Burke, Kristian Aquilina, Ulrike Löbel, Isidro Cortes-Ciriano, Darren Hargrave, Mette Jorgensen, Flora A. Jessop, Tim H. H. Coorens, Adrienne M. Flanagan, Kieren Allinson, Inigo Martincorena, Thomas S. Jacques, Sam Behjati

## Abstract

Cancer predisposition syndromes mediated by recessive cancer genes generate tumours via somatic variants (second hits) in the unaffected allele. Second hits may or may not be sufficient for neoplastic transformation. Here, we performed whole genome and exome sequencing on 479 tissue biopsies from a child with neurofibromatosis type 1, a multi-system cancer-predisposing syndrome mediated by constitutive monoallelic *NF1* inactivation. We identified multiple independent *NF1* driver variants in histologically normal tissues, but not in 610 biopsies from two non-predisposed children. We corroborated this finding using targeted duplex sequencing, including a further nine adults with the same syndrome. Overall, truncating *NF1* mutations were under positive selection in normal tissues from individuals with neurofibromatosis type 1. We demonstrate that normal tissues in neurofibromatosis type 1 commonly harbour second hits in *NF1*, the extent and pattern of which may underpin the syndrome’s cancer phenotype.

## MAIN TEXT

In recessive tumour predisposition syndromes, one allele is mutated in the zygote (or, rarely, in early embryogenesis) whilst the second allele is inactivated by subsequent somatic mutation (second hit) (**Fig. 1a**). Although a second hit would ordinarily be expected to lead to neoplasia, it is possible that some cells remain phenotypically normal in the presence of biallelic mutation, just as oncogenic mutations have been reported within healthy tissues. Recent studies of normal adult tissues have revealed bonafide cancer-causing (driver) mutations that accumulate with age and exposure to environmental mutagens, primarily in exposed epithelial tissues^1–4^. The acquisition of mutations in normal tissues may be accelerated by germline mutations perturbing the fidelity of DNA replication, as seen in normal intestinal crypts of patients with a mutant DNA polymerase^5^. Furthermore, in children with malignant rhabdoid tumours (cancers driven by biallelic inactivation of *SMARCB1*) we have observed normal tissues that share a genetic ancestor with the nearby tumour and harbour the same somatic *SMARCB1* hit, without an elevated mutation rate^6^ (**Fig. 1b**). We therefore speculated that second hits may occur in normal tissues of predisposed individuals that are unrelated to tumour lineages or affected by hypermutation, which we set out to investigate here (**Fig. 1c**).

**Fig. 1:**
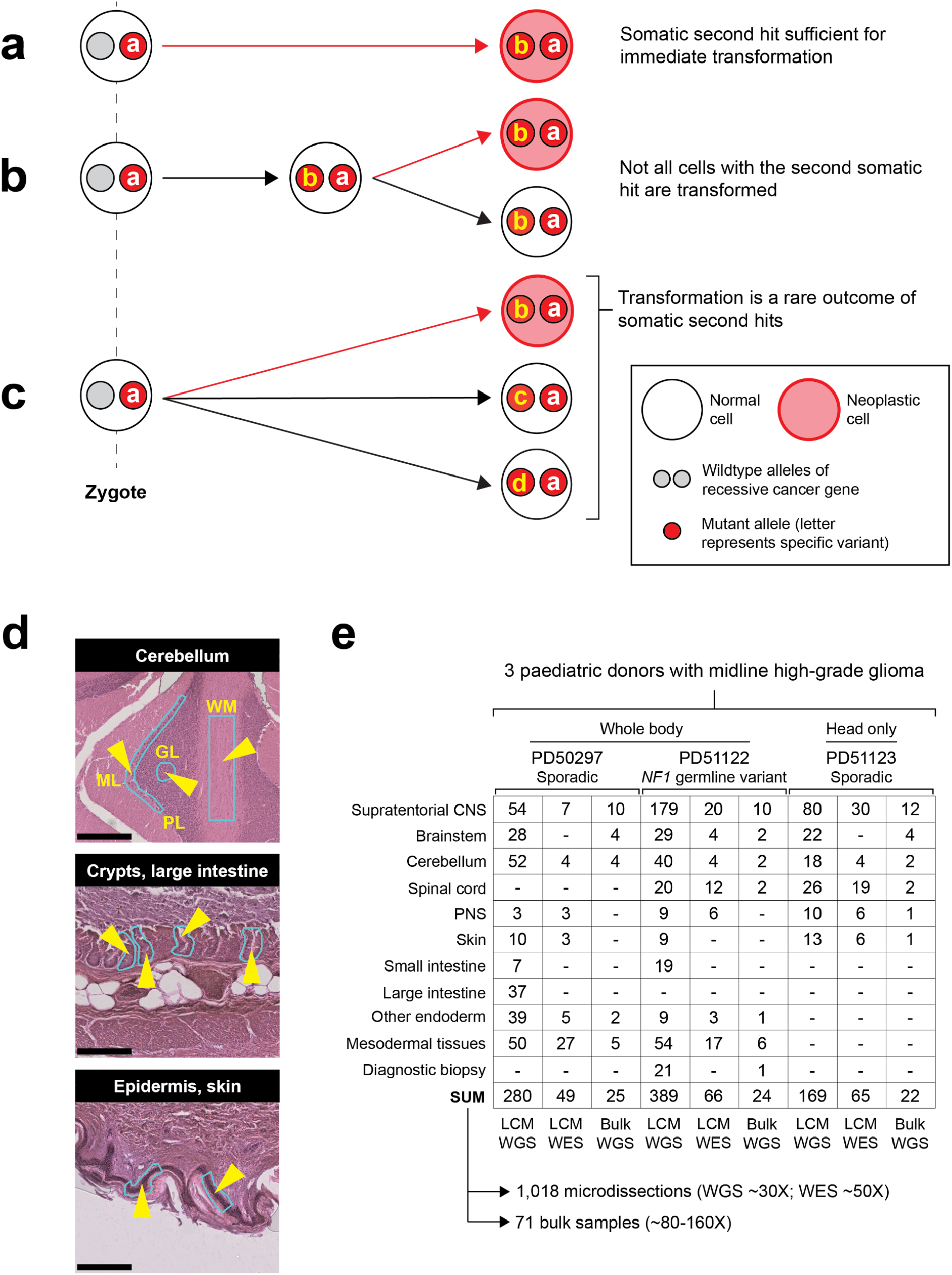
**Concepts and experimental approach. a**, The second mutation in a recessive predisposition syndrome is typically thought to lead to neoplasia. **b**, Some second hits may be found in the adjacent normal tissue to a childhood cancer, indicating that their presence is insufficient for neoplastic transformation. **c**, The possibility remains that second hits may be sustained in normal tissues that are independent of the cancer cell lineage. **d**, Histological images of three illustrative microdissected tissues are shown. The layers of the cerebellar cortex are annotated on the uppermost image: ML, molecular layer; PL, Purkinje layer; GL, Granular layer; WM, white matter. The light blue outlines with yellow arrowheads on the images are representative regions microdissected. The scale bars represent 500, 250 and 250 µm (top to bottom). **e**, Experiment overview, detailing the number of bulk and laser capture microdissection (LCM) sequences generated per anatomical region per child. Note that this includes all biopsies, irrespective of tumour involvement. A similar table, limited to the tumour biopsies used in the high grade glioma driver mutation identification, is provided as Supplementary Table 2. CNS, central nervous system; PNS, peripheral nervous system.

Neurofibromatosis type 1 is a complex multi-system disorder that predisposes to neoplasia. It is caused by germline mutation in the *NF1* gene, a tumour suppressor gene that encodes neurofibromin, a negative regulator of intracellular RAS/MAPK signalling. The neoplastic phenotype of *NF1* is variable and tends to affect neuroectodermal lineages, although tissues derived from other germ layers also have an increased risk of cancer. An essential diagnostic feature of neurofibromatosis is the café au lait spot, a macroscopically visible clonal expansion of melanocytes^7,8^. Other neoplastic manifestations, which exhibit variable penetrance, include neurofibromas, skeletal dysplasias, leukaemias, malignant peripheral nerve sheath tumours and gliomas^9–12^. In all these lesions, *NF1*, as a recessive cancer gene, exhibits a second mutation, not infrequently as the sole detected somatic driver event^11,12^, consistent with Knudson’s two hit hypothesis^13^.

We performed a post mortem study of three children aged <10 years old with high grade midline gliomas: two (PD50297, PD51123) with sporadic tumours (*H3F3A* K27M mutant) and one with neurofibromatosis type 1 with a pathogenic truncating *NF1* (c.3113+1G>A) germline mutation. Our key question was whether normal tissues across the body harboured driver events, in particular in the predisposed child. We extensively sampled normal tissues and neoplasms (**Supplementary Table 1**), which, in the case of the child with neurofibromatosis, included the brain tumour, a subcutaneous spindle cell lesion (**Supplementary** Fig. 1), and a café au lait spot. Guided by parental wishes, we surveyed central nervous system (CNS) tissues in all three children and extracranial tissues in two of them, including the predisposed child. None of the children had been pre-treated with cytotoxic chemotherapy. Radiotherapy was given to the two children with sporadic tumours.

In total, we performed whole genome sequencing (WGS) on 838 microdissected groups of cells (median coverage 28×), using an established approach that we and others have pursued in the study of normal tissue genomes^14^ (**Figs. 1d, e**). We supplemented this with additional bulk tissue WGS (n = 71) and whole exome sequencing (WES) of other microdissected tissues (n = 180) (**Fig. 1e**). We assembled catalogues of all classes of mutations (substitutions, indels, rearrangements and copy number changes (**Supplementary Tables 3**-9)) using a validated variant calling pipeline (**Methods**). To exclude low-level tumour contamination of normal tissues, we quantified the extent of tumour infiltration in each sample (including samples distant from the tumour) by searching for the mutations assigned to the tumour’s phylogenetic trunk (**Methods**, **Supplementary Tables 3**, 10).

We identified somatic driver variants in both neoplastic and normal tissues (**Table 1**, **Supplementary Tables 11**-15). Gliomas exhibited a multitude of driver mutations in cancer genes known to operate in gliomagenesis^15,16^. The normal tissues of the two non-predisposed children bore comparatively few cancer-associated mutations, yielding only a *CREBBP* frameshift mutation (p.Q2199fs*99) within a single colonic crypt (PD50297g_lo0012) by our standard pipeline. Further inspection of the copy number data revealed one more putative driver, chromosome 11p loss of heterozygosity (LOH) (**Supplementary** Fig. 2) in a nerve (PD51123t_lo0028), a variant commonly reported in Wilms tumour, rhabdomyosarcoma and hepatoblastoma^17^.

**Table 1:**
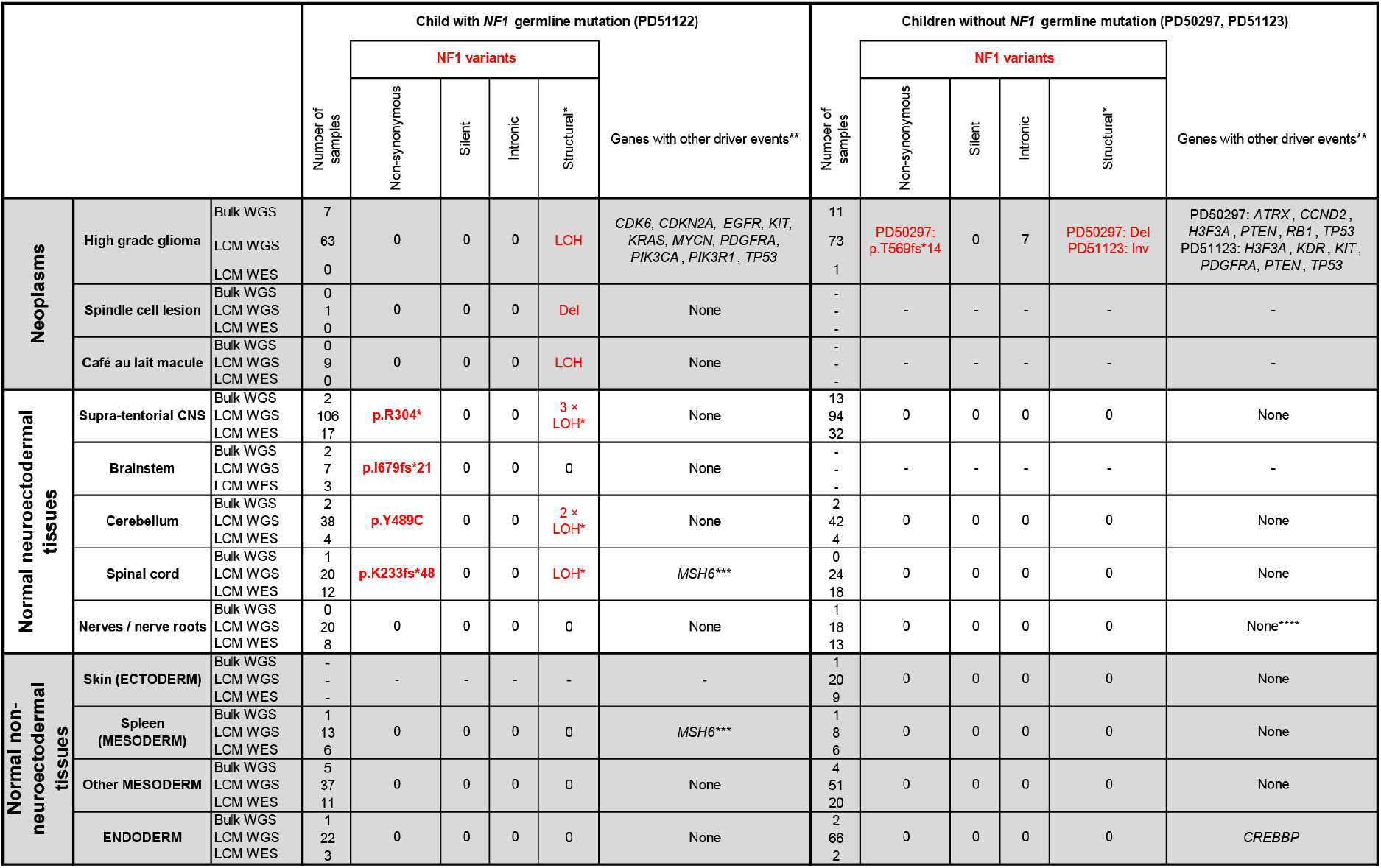
***NF1* mutations and driver events identified by whole genome or exome sequencing of bulk tissues or microdissections.** Individual driver variants can be found in the lists of mutations provided in Supplementary Tables 11-15. * Identification of LOH events was only possible because we could definitively phase parental alleles (**Methods**). The low cell fraction of many of these meant that it was not possible to determine the breakpoint. When counting LOH events, those called from sequences derived from the same original bulk biopsy are treated as the same event and those from different biopsies as unique. These events should not be considered when comparing mutations against the other two children because their alleles over chromosome 17 could not be phased. ** In sequences from the high grade glioma, we only considered mutations in genes recognised in a large meta-analysis to be drivers of these neoplasms (**Methods**). ***One additional *MSH6* frameshift mutation (p.F1088fs*2) was noted in the spinal cord (PD51122v_lo0008) and spleen (PD51122z_lo0017) of the child with neurofibromatosis type 1; however, this remained heterozygous without evidence of hypermutation and did not co-occur with somatic *NF1* mutation, making this of uncertain significance. ****Chromosome 11p LOH was identified in a single sample after manual inspection of the copy number output although it was too low fraction to be detected by the copy number caller. LOH, loss of heterozygosity; Del, deletion; Inv, inversion; CNS, central nervous system.

By contrast, in the child with neurofibromatosis type 1, we found bonafide somatic *NF1* driver point mutations (**Table 1)**, that either truncated the gene (p.R304*, p.K233fs*48, p.I679fs*21) or were likely oncogenic based on recurrence, *in silico* predictions, correlation between genotype and phenotype, and functional studies (p.Y489C)^18,19^. These mutations were detected by a combination of whole genome and whole exome sequencing of microdissections or bulk tissues. They occurred in anatomically distant regions of the central nervous system (left parietal cortex, cerebellar hemisphere, or spinal cord). The affected tissues appeared macroscopically and microscopically normal, and were not correlated with focal areas of signal intensity on imaging (a feature often found in the brains of children with neurofibromatosis type 1^20^) (**Supplementary** Figs. 3-4). The variant allele frequencies (VAFs) of second *NF1* hits indicated clone sizes as large as 56% of cells (clone size = 2 x VAF) in a microdissection (hundreds of cells) and 19% in a bulk tissue (a macroscopic piece of tissue). *NF1* second hits were independent of those found in the clonal lesions (glioma, spindle cell lesion, café au lait spot; **Fig. 2a**). In particular, as both copies of *NF1* in the glioma were already inactivated, there should be no selection pressure for further loss-of-function mutations in *NF1* in tumour cells. Given this and our ability to detect tumour contamination accurately (**Methods**), the *NF1* mutations in normal tissues are not the result of tumour cells infiltrating normal tissues. The mutation burden of *NF1* null normal tissues was inconsistent with a recent clonal expansion (**Fig. 2b, Methods**). Like the spindle cell lesion and café au lait spot, no additional driver mutations were identified in the *NF1* null histologically normal tissues.

**Fig. 2:**
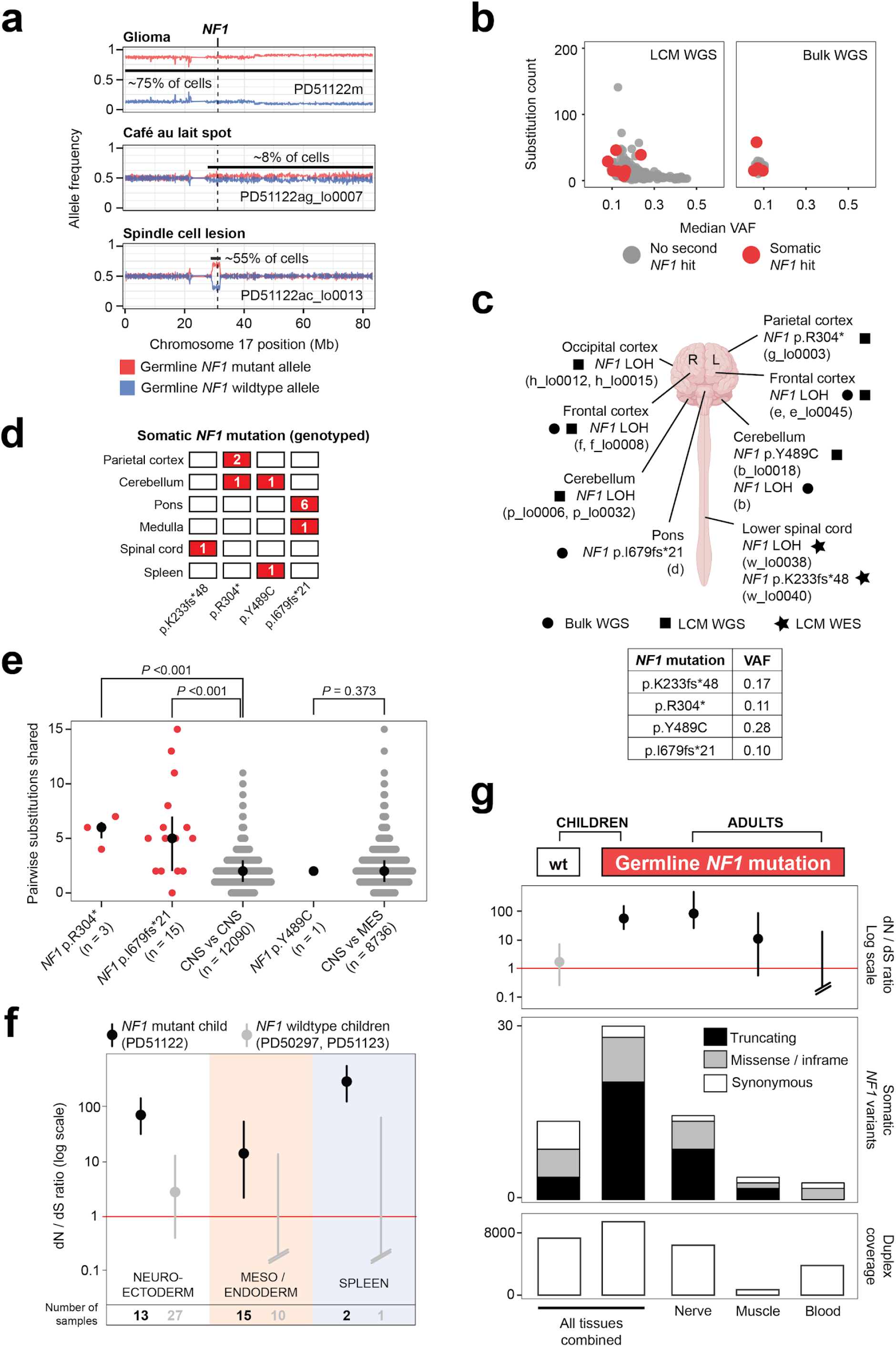
**NF1 null clones pervade the normal tissues of individuals with Neurofibromatosis type 1. a**, Loss of the wildtype *NF1* allele is an independent event in each of the three neoplasms in PD51122. The allele frequency on the Y-axis represents a rolling window of 50 SNPs. **b**, The substitution count and median VAF of normal brain tissues, colour-coded by the presence or absence of biallelic *NF1* mutation. **c**, Schematic of the brain and spinal cord, outlining the location of each somatic *NF1* mutation in normal tissue from PD51122, prior to genotyping the point mutations, discovered by different sequencing methods. The VAF of the point mutations in the affected tissue is shown in the table beneath, rounded to two significant figures. Image created with BioRender.com. R, right; L, left. **d**, Distribution of somatic *NF1* point mutations across tissues from PD51122 (above the locus-specific error rate, **Methods**). If a mutation is found within a tissue, the corresponding box is coloured red. The number within the box indicates the number of samples from that tissue that are affected. **e**, Pairwise comparison of the number of substitutions shared between normal tissue biopsies, according to whether or not they possess the same *NF1* second hit. The black line represents the interquartile range and the black dot is the median value. *P* values were generated using one-sided permutation tests (Methods). Note that “n” refers to the number of pairwise comparisons, not samples. CNS, central nervous system; MES, mesoderm. The CNS and MES groups exclude normal tissues with a second *NF1* hit. **f**, dN/dS ratios for truncating variants, according to germ layer and *NF1* germline mutation status. The dot represents the maximum likelihood estimate and the lines represent the 95% credible interval. When the lower bound of the credible interval is above 1 (red line), there is statistically significant positive selection. Credible intervals falling below the boundary of the plot are terminated with slanted double lines. **g**, Normal tissues from adults with neurofibromatosis type 1 are grouped by tissue type and evaluated for an excess of non-synonymous variants in *NF1*, and compared with the three index children. Upper, dN/dS ratios for truncating mutations; middle, counts of variants in *NF1*; lower, total duplex coverage (**Methods**) over *NF1* in each group.

To expand the breadth and depth of *NF1* mutations detected within the normal tissues of the predisposed child, we employed two strategies. First, we re-examined all the child’s sequencing data for evidence of LOH of the *NF1* locus (chromosome 17q). As this child’s glioma harboured complete LOH of chromosome 17 (**Fig. 2a**), we were able to increase the sensitivity to detect allelic shift by obtaining definitive chromosome 17 haplotypes from phasing allele-specific single nucleotide polymorphisms (**Methods**, **Supplementary Tables 3-4**, **Supplementary** Fig. 5). This haplotype-resolved copy number calling revealed six instances of LOH in normal tissues (**Table 1**), of which at least two were demarcated by distinct breakpoints consistent with independent events (**Supplementary** Fig. 5). We captured LOH-driven *NF1* null clone sizes as small as 2% (PD51122b, cerebellum) and up to 13% (PD51122h_lo0012, occipital cortex), and no second *NF1* hit was related to a neoplasm. Curiously, while this approach did not yield any *NF1* null clones in any non-neuroectodermal lineage, it did identify rare examples where the germline mutant allele was lost in microdissections of tissues derived from other germ layers (bladder muscle (PD51122s_lo0012) and a renal tubule (PD51122u_lo0009)) (**Supplementary** Fig. 6). All *NF1* second hits identified thus far are shown in Figure 2c.

Second, to deepen our search for *NF1* null normal cells, we assessed all samples for evidence of *NF1* point mutations that had been previously called in normal tissues. Three variants (p.R304*, p.I679fs*21, p.Y489C) were detectable in more than one biopsy above the locus-specific error rate (**Methods**, **Fig. 2d**, **Supplementary Table 16**). There were two possible explanations: either the shared variants arose prior to seeding of the anatomical areas in which the mutations were found, or mutations appeared independently in different tissues. We could establish which scenario was more likely by comparing the total number of mutations across the genome that were shared between affected tissues: those with a common developmental root would possess more. For *NF1* mutations that spanned only regions of the brain (p.R304*, p.I679fs*21), affected tissues shared significantly more mutations with each other than unaffected brain regions did (*P* <0.001 for both mutations, permutation test; **Fig. 2e**, **Methods**), implicating a common ancestor in their development, although convergent evolution with shared selection pressures between developmentally-related tissues is also possible. The same could not be said for the *NF1* mutation found in both the brain and spleen when compared to normal tissues of the CNS and mesoderm (p.Y489C; *P* = 0.373, permutation test; **Fig. 2d**), meaning that they likely developed independently.

Taken together, the multiple lines of enquiry we had pursued thus far pointed towards an enrichment of *NF1* non-synonymous mutations within the normal tissues of a predisposed individual but not wildtype individuals, at least within the brain and spinal cord. It seems likely that this pattern of *NF1* mutation has emerged as a consequence of positive selective pressure, given the absence of the concomitant silent and intronic *NF1* variants (**Table 1**) that would be expected under a neutral model.

To establish statistically whether there is positive selection for non-synonymous variants in *NF1* in predisposed normal tissues, we re-interrogated 60 normal tissue samples (21 from the predisposed child and 39 from the unaffected children) by duplex sequencing of the *NF1* gene^21^. Duplex sequencing, through barcode tagging of both strands of DNA molecules, enables highly sensitive and specific mutation calling which may deliver sufficient variants for formal statistical assessment of selection through the non-synonymous to synonymous variant ratio (dN/dS)^15^. Duplex sequencing delivered a total of 29 non-synonymous (21 truncating; eight missense or inframe) and two synonymous *NF1* mutations in the predisposed child. By contrast, in the normal tissues of the two children without neurofibromatosis we detected nine non-synonymous (four truncating; five missense or inframe) and five synonymous *NF1* mutations. Calculation of the dN/dS ratio provided strong evidence of positive selection for truncating *NF1* variants in normal tissues from the child with a germline predisposition (**Fig. 2f**). Interestingly, the spleen had a particularly high proportion of truncating variants, and, when analysed separately from other tissues, it had the highest dN/dS ratio (**Fig. 2f**). This finding was of interest given that neurofibromatosis type 1 predisposes to juvenile myelomonocytic leukaemia, which always (as per the diagnostic definition) involves the spleen^22^, whether through entrapment of leukaemic cells or as their organ of origin.

Next, we extended our analysis into normal tissue from adults with neurofibromatosis type 1 (**Supplementary Table 17**). We were able to obtain normal peripheral nerves, muscle tissue, or blood from nine individuals who underwent extensive surgical resections for sarcoma. The principal cellular material of peripheral nerves is made up of Schwann cells which are derived from the neuroectoderm, whereas muscle and blood develop from mesoderm. Consistent with the pattern of mutation we observed in paediatric tissues, we found, by duplex sequencing, a stark excess of truncating *NF1* mutations in peripheral nerves, indicative of positive selection (**Fig. 2g**).

In this study of *NF1* mutations in individuals with neurofibromatosis type 1, we observed independent second *NF1* hits in macroscopically and histologically normal paediatric and adult tissues. Multiple lines of evidence arrive at the same conclusion: in neurofibromatosis type 1, non-synonymous second somatic mutations of *NF1* are selected for in histologically normal tissues. Although *NF1* is a ubiquitously expressed gene, the tissue distribution of neoplasms associated with neurofibromatosis type 1 is not random, showing a predilection for neuroectodermal lineages^23^ which is mirrored in the distribution of second *NF1* hits we identified. Unlike our study though, where these mutations pervaded the CNS, most brain tumours that arise in children with neurofibromatosis type 1 are localised to the optic pathway and brainstem^23,24^. Our findings may thus explain some, but not all, of the cancer phenotype associated with neurofibromatosis type 1.

Three factors, unrelated to the germline mutation status of *NF1*, may also have contributed to the number of non-synonymous mutations we observed. The first is age, as neutral mutations accrue with time, and the second is the size of the *NF1* gene. *NF1* has one of the largest footprints of any gene (8520bp coding sequence compared to a median length across human genes of 1257bp^25^), meaning there are simply more sites to mutate. Hypermutation is a third factor that might augment the rate of driver mutation acquisition, but our comprehensive study of the index case found no evidence to support that here. Importantly, all these factors are accounted for by our model of selection (dN/dS), meaning that they cannot explain our data. We can assume, then, that the strength of the signal we see in the carriers compared to unaffected children is the result of selective pressure specifically for second hits. The fact that this is so readily apparent in the extensively studied child suggests that this occurs from an early age, possibly even during development (**Figs. 2b, e**). Given the extent to which we observed *NF1* loss-of-function variants in normal tissues, it seems reasonable to propose that even though in certain contexts second hits may sufficient to cause neoplasia^10,11^, as suggested by our case’s café au lait spot and spindle cell lesion, transformation to a discernible tumour is an uncommon immediate outcome of biallelic *NF1* loss.

This finding may represent a fundamental principle of *NF1* mutation in neurofibromatosis type 1, the precise nuances and clinical implications of which will require extensive surveys in human tissues^26^. Consistent with our data, mouse models support a complex relationship between cellular genotype and phenotype in neurofibromatosis type 1: the genetic background of the mouse, the identity of the cell in which *NF1* is inactivated, the presence of cooperating somatic mutations, and the status of *NF1* function in neighbouring cells, all appear to affect tumour development^27,28^. From a practical, clinical point of view it is conceivable that the extent of second *NF1* hits in normal tissues represents a quantifiable link between germline genotype and cancer risk. In the broader context of recessive cancer predispositions, our findings call for systematic investigations to establish whether second hits occur commonly in such predispositions or delineate a particular group of syndromes.

## METHODS

### Sample collection

Study of the discovery cohort of three children was approved by NHS research ethics committees (PD50297 - HRA East Midlands Derby REC, 08/H0405/22+5; PD51122 & PD51123 - London Brent REC, 16/LO/0960). Each autopsy was undertaken at the child’s local neuropathology unit, with parental consent, 1-7 days following death. Samples were snap frozen at the point of sampling with adjacent brain tissue taken for immediate formalin fixation and processing in the local diagnostic laboratory. A full list of the samples taken is provided in Supplementary table 1.

Study of the validation cohort of adults with neurofibromatosis type 1 was approved by NHS research ethics committees (20/YH/0088, IRAS 272816). Patients with a diagnosis of neurofibromatosis type 1 who were undergoing a resection for sarcoma consented to the use in research of normal tissue removed as part of the resection but distant from the lesion, or of blood samples. Solid tissue samples were immediately frozen in liquid nitrogen. Blood samples were centrifuged and plasma and cellular fractions separated prior to freezing.

### Preparation of samples for sequencing - tissue processing and DNA extraction

A subset of the bulk samples in the discovery cohort underwent bulk DNA extraction using either the DNeasy Blood & Tissue Kit (Qiagen, Hilden, Germany), AllPrep DNA/RNA Mini Kit (Qiagen) or the Gentra Puregene Blood Kit (Qiagen). The choice of samples to undergo bulk DNA extraction was guided by prior understanding of the clonal architecture of the tissue. For example, intestinal biopsies were not subject to bulk DNA extraction. This is because their clonality meant that pseudo-single cell genome readouts, rather than a single polyclonal amalgamation, could be achieved by microdissection of individual crypts instead^2^. Similarly, bulk DNA extraction was not performed for samples taken from the interface of tumour and normal as it was hoped microdissection would better isolate the tumour and normal tissue compartments.

The remaining tissue from solid organ biopsies in the discovery cohort was fixed in PAXgene (PreAnalytiX, Hombrechtikon, Switzerland), according to the manufacturer’s instructions, and processed in preparation for laser capture microdissection using an established protocol^14^. To ensure correct feature labelling of nervous system structures during microdissection, reference slides were generated using 4-micron thick sections mounted on SuperFrost Plus slides (VWR International, Lutterworth, UK) and reviewed by the neuropathologist who performed the autopsy. The sections subject to microdissection were 16-microns thick and mounted on polyethylene naphthalate membrane slides (Leica Microsystems, Wetzlar, Germany). The microdissected tissue then underwent lysis and further DNA extraction^14^.

DNA was extracted from solid tissues from the validation cohort using the DNeasy Blood & Tissue Kit (Qiagen), and from blood using the Gentra Puregene Blood Kit (Qiagen).

### Preparation of samples for sequencing - library preparation for whole genome and whole exome sequencing

In the discovery cohort, the NEBNext Ultra II DNA Library Prep Kit (New England Biolabs, Ipswich, Massachusetts, USA) was used for preparation of DNA extracted from the bulk samples whilst the protocol for microdissected tissue used the NEBNext Ultra II DNA FS Library Prep Kit (New England Biolabs) instead. For the microdissected libraries subject to whole exome sequencing, the SureSelect Human All Exon V5 bait set (Agilent Technologies, Santa Clara, California, USA) was used. A full list of the successfully generated whole genome and exome sequences can be found in Supplementary tables 3 and 4 respectively.

### Preparation of samples for sequencing - library preparation for duplex sequencing of the *NF1* gene

DNA was sheared to an average size of 450bp by focused ultrasonication using the Covaris 644 LE220 instrument (Covaris, Woburn, Massachusetts, USA) in 120uL. It was purified using a 0.8X soluble-phase reversible immobilisation (SPRI) bead ratio and eluted in 30uL nuclease-free water (NFW). DNA fragments were blunted in a final reaction volume of 30 μl including 3uL 10X Mung Bean Buffer (Cat. #2420A; Takara Bio Inc., Kusatsu, Japan), 0.125uL Mung Bean Nuclease (Cat. #2420A; Takara Bio Inc.), 1.875uL of NFW and 25uL DNA. The reaction was incubated at 37°C for 10 min with the lid tracking 5°C above. Samples were purified using 2.5X SPRI beads and eluted in 15uL NFW. 10 μl was used as input into an A-tailing reaction, containing 1.5uL T4 DNA ligase buffer (Cat. #B0202S; New England Biolabs), 1.5uL 1mM dATP/ddBTP (Cat. #N0440S; New England Biolabs / Cat. #27204501; GE HealthCare, Chicago, Illinois, USA), 1.5 μl Klenow fragment (3′ to 5′ exo-, Cat. #M0212L; New England Biolabs) and 0.5 μl T4 Polynucleotide Kinase (Cat. #M0201L; New England Biolabs). The reaction was incubated at 37°C for 30 min with the lid tracking 15°C above. The whole sample of 15μl was taken into the ligation reaction mix, which consisted of 30uL UltraII Ligation MM (Cat. #E7595L; New England Biolabs), 1uL UltraII ligation enhancer (Cat. #E7595L; New England Biolabs), 1.25 μl xGen Duplex Seq Adapters (Cat. #1080799; Integrated DNA Technologies, Coralville, Iowa, USA) and 12.75 μl NFW. The reaction was incubated at 20°C for 20 min, with the lid temperature off. Ligated DNA was cleaned up using SPRI beads and eluted in 40uL NFW.

DNA was quantified by qPCR using a KAPA library quantification kit (Cat. #KK4835; Kapa Biosystems, Wilmington, Massachusetts, USA). The supplied primer premix was first added to the supplied KAPA SYBR FAST master mix. In addition, 20 μl of 100 μM NanoqPCR1 primer (HPLC, 5′-ACACTCTTTCCCTACACGAC-3′) and 20 μl of 100 μM NanoqPCR2 primer (HPLC, 5′-GTGACTGGAGTTCAGACGTG-3′) were added to the KAPA SYBR FAST master mix. Samples were diluted 1:500 using NFW and reactions were set up in a 10μl reaction volume (6μl master mix, 2μl sample/standard, 2μl water) in a 384 well plate. Samples were run on the Roche 480 Lightcycler and analysed using absolute quantification (second derivative maximum method) with the high sensitivity algorithm. The concentration (nM (fmol/μl)) was determined as follows: mean of sample concentration × dilution factor (500) × 452/573/1,000 (where 452 is the size of the standard in bp and 573 is the proxy for the average fragment length of the library in bp), and multiplied by an adjustment factor of 1.5. Samples were diluted to the desired fmol amount in 25μl using NFW.

Libraries were subsequently PCR-amplified in a 50-μl reaction volume comprising 25μl of sample, 25μl NEBNext Ultra II Q5 Master Mix and unique dual index (UDI) containing PCR primers (dried). The reaction was cycled as follows: step 1, 98°C 30 s; step 2, 98°C 10 s; step 3, 65°C 75 s; step 4, return to step 2 13 times; step 5, 65°C for 5 min; step 6, hold at 4°C. The number of PCR cycles is dependent on the input:

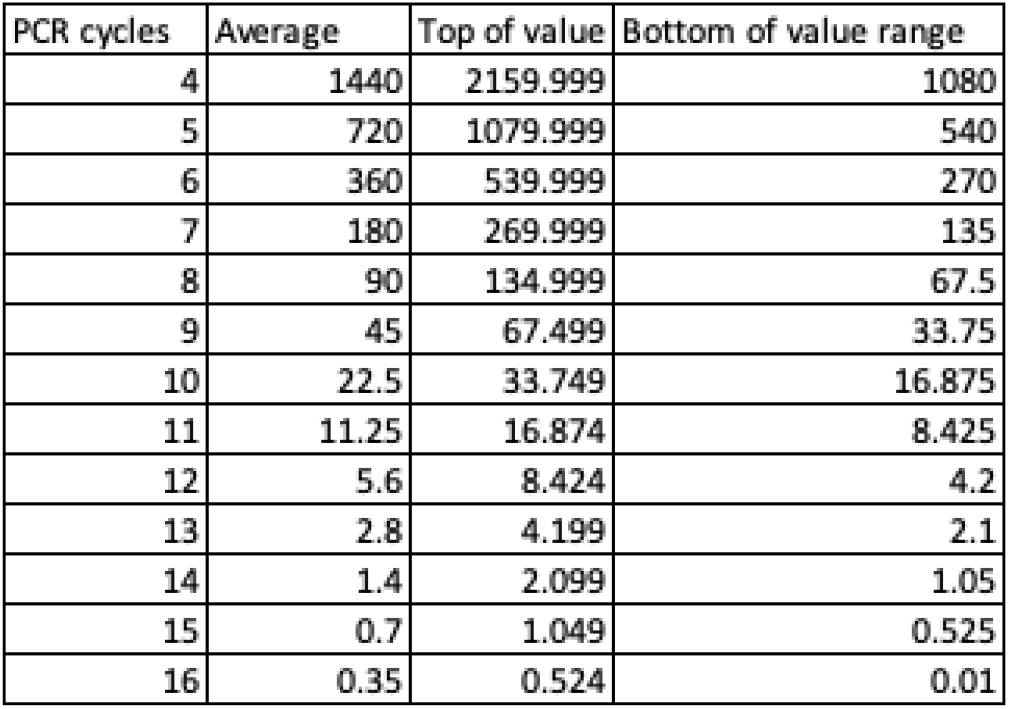

The PCR product was subsequently cleaned up using two consecutive 0.7× AMPure XP clean-ups. Each sample was quantified using the AccuClear Ultra High Sensitivity dsDNA Quantification kit

(Biotium, Fremont, California, USA). Hybrid capture was performed using TWIST hybe reagents. Samples were pooled for hybridisation with 1-4ug of PCR-amplified material per capture reaction.

### DNA sequencing

All DNA sequences were generated on the Illumina NovaSeq sequencing platform, generating paired-end 150bp sequences. Sequences were aligned to the GRCh38 human reference genome using the Burrows-Wheeler Aligner (BWA-MEM)^29^.

### Assessment of DNA sequencing data quality

The CollectWgsMetrics tool from Picard (version 2.26.10) was used on the whole genome sequencing data to determine the median coverage and duplicate rate. The coverage cap was set to 500. CollectHsMetrics, also from Picard, was deployed for the whole exome sequencing data to determine the on-target coverage and duplicate rate.

### Assessment of sample-to-sample concordance

Conpair (version 0.2)^30^ was run between every sample that underwent whole genome or whole exome sequencing and a designated matched normal sample (PD50297b, PD51122q, PD51122aa3). Two samples were found to have >0.5% contamination and were excluded from the final data set analysed here (PD51122t_lo0011, PD51122t_lo0012). The remaining samples all showed a concordance of >99.5% with their matched normal.

### Variant calling and filtering for whole genome and whole exome sequencing - substitutions

Substitutions were called using the CaVEMan algorithm (version 1.15.1)^31^, unmatched against an *in silico* normal sample (PDv38is_wgs) for the whole genome data and matched (blood or skin) for the whole exome data. All called variants were initially filtered using soft flag filters contained within the VCF (ASRD ≥0.87 & CLPM = 0) to eliminate BWA-MEM-associated mapping errors. Further artefacts introduced as a result of cruciform DNA formation during DNA library preparation, a recognised consequence of the low input method used for microdissected tissue DNA in particular, were removed using previously described filters^14^. Briefly, the artefacts generated by this phenomenon can be identified by their tendency to occur at similar alignment start positions on supporting reads.

The extended nucleotide contexts that passed variants were found in was examined. Some sequences derived from microdissected tissues showed an abundance of substitutions at loci with an upstream GGGTGGTCT, as has been previously noted^5^. Further hotspots were identified at loci with a preceding AGACCA and in regions of long (≥9bp) A or T repeats. Any substitution called where at least 8 of the upstream 9 bases matched GGGTGGTCT, or where the preceding 6 bases were AGACCA, or which occurred upstream or downstream of a poly-A or poly-T repeat ≥9bp long, was excluded. For the whole exome sequencing data, which was generated solely to expand our survey of the driver landscape, the only additional filter implemented was a cutoff of at least four supporting reads to consider a variant during driver curation.

Common SNPs were removed from the remaining unmatched whole genome sequencing by eliminating called at loci reported to be mutated in gnomAD (version 3.1.1) with an allele frequency >0.1%. Recurrent artefacts identified within the Panel of Normals mask created by the Brain Somatic Mosaicism Network were also removed (PON.q20q20.05.5.fa). Bidirectional read support was required for passed substitutions to be considered further. At least one supporting read needed to be found in both directions and, where total locus coverage was >10, at least 15% of supporting reads needed to be found in both directions.

Genotyping of the unique, remaining mutations across all samples from a single case was performed using pileup (https://github.com/cancerit/vafCorrect, version 5.6.0), with defined mapping quality (MQ 30) and base quality (BQ 25) thresholds. Variants found at loci with a universally low coverage (≤10) across all case samples were removed, as were those which were not supported by at least four mutant reads in one or more samples. Examining the genotypes of variants across all non-CNS samples, where the risk of tumour infiltration was remote, remaining germline variants were excluded through the use of a one-sided exact binomial test. This test assumed that the fraction of reads supporting a germline mutation were drawn from a binomial distribution where *P* = 0.5, or 0.95 in the case of male sex chromosomes (null hypothesis). Somatic mutations should exist at a fraction beneath these cutoffs (alternative hypothesis). The Benjamini-Hochberg multiple hypothesis testing correction was applied to the resultant *P* values with a cutoff of *q* < 10^-5^, as has been done before^32,33^.

Certain artefacts, or germline mutations on DNA segments with constitutional copy number variation, may appear in multiple clonal samples at a low allele fraction. These calls were filtered by fitting a beta binomial distribution to the mutant read depth and total depth of each variant shared across two or more microdissected regions of epithelium and removing variants whose rho (overdispersion parameter) value was <0.1^32,33^. This was limited to epithelial samples as they should be tumour-free but sufficiently clonal to minimise the loss of true mosaic variants from within polyclonal tissues through their apparent underdispersion.

Substitutions that were identified across multiple cases and were not bonafide driver mutations likely represent recurrent artefacts, often as a result of mismapping. They were removed and only variants where >90% of reads at the locus had MQ >30 were kept. Any variants found within 5bp of an indel call within the unfiltered Pindel file (used for indel calling, see below) were also excluded as possible mapping artefacts. The final substitution filtering step involved calculating the background sequencing error rate for each mutation found in one of the three cases using the sequencing data from the other two cases, similar to the method employed by Shearwater^1,32,33^. Again, rare, shared driver events were not subject to this step. Variants that exceeded the error rate modelled from both unrelated bulk sequencing and microdissected samples were kept for subsequent analysis.

### Variant calling and filtering for whole genome and whole exome sequencing - small insertions and deletions (indels)

For indels, the Pindel algorithm (cgpPindel version 3.5.0)^34^ was used and 1bp indels at homopolymer runs of 9bp or longer were removed. As with the substitutions, the whole exome sequencing data required a minimum variant depth: this was set to five for indels. No further filtering was performed for the whole exome data before driver curation.

The same read direction, mean coverage, binomial test, beta binomial distribution and locus MQ (>90% MQ >30) filters were applied to the indels from the whole genome sequencing data as for the substitutions. The only notable differences were that a variant needed to have five or more supporting reads in at least one sample and the rho value cutoff was set to 0.2. For indels, genotyping was carried out using Exonerate (https://github.com/cancerit/vafCorrect, version 5.6.0) which may annotate reads as “ambiguous” or “unknown” if they cannot be assigned to wildtype or the interrogated mutant allele. If >10% of reads at a locus were assigned to these categories, the variant was discarded.

### Variant calling and filtering for whole genome and whole exome sequencing - copy number changes

Somatic copy number variants in the whole genome data were identified by two different callers. The first, Battenberg (cgpBattenberg version 3.5.3)^35^, was run using the bulk data only with bulk blood or skin samples acting as the matched normal sample. The second, ASCAT (AscatNGS version 4.3.2)^36^, was applied to all samples using the same matched normal samples and a segmentation penalty of 100. To eliminate technical sources of variation in coverage that add noise to the logR profile, ascatPCA was applied (https://github.com/hj6-sanger/ascatPCA). This performs a principal component analysis using a panel of unrelated, normal samples to define recurrent regions of artefact change and uses these to eliminate false positive calls^37^. The quality of the resultant copy number calls could be assessed with a “goodness of fit” metric. Furthermore, orthogonal validation was also possible in the samples with a high substitution burden, especially if they had a high tumour purity. This is because the VAF of substitutions on a segment must be concordant with its copy number state and the purity of the sample they are found in. To this end, we ran CNAqc (version 1.0.0)^38^ using the substitution calls and the ascatPCA copy number calls and purity estimates. Using the original tumour purity estimate from ascatPCA, we found that only 30% (26/88) of PD51122 sample copy number calls with ≥40% tumour purity passed the CNAqc check using default parameters, compared to 92% (48/52) and 89% (34/38) of PD50297 and PD51123 tumour samples respectively. On closer inspection, whilst the purity estimates between ascatPCA and our method of estimating tumour purity (see ‘Estimating Tumour Infiltration’ below) were very similar for the near-diploid tumours, ascatPCA provided higher than expected purity estimates for the aneuploid PD51122 (**Supplementary figure 7A**). Our method of estimating purity showed good concordance with Battenberg calls in all tumours for the bulk samples (**Supplementary figure 7B**) so we ran CNAqc again for the PD51122 tumour samples with the purity from our method. This markedly improved the concordance between substitutions and copy number calls with 76% passing (67/88), indicating our method of inferring purity better accounted for the copy number calls and substitution VAFs. In Supplementary table 3, the “PASS/FAIL” CNAqc column uses the original ascatPCA purity except for PD51122 where the ≥40% tumour purity samples use the purity derived from our approach. We kept copy number data from all samples where the ascatPCA goodness of fit was ≥95% and tumour purity (our method) <40%. For samples with ≥40% tumour purity (our method), we kept calls if the average number of reads per chromosome copy was ≥5 and either ascatPCA goodness of fit was ≥95% or the sample passed CNAqc.

### Variant calling and filtering for whole genome and whole exome sequencing - structural variants

GRIDSS (version 2.9.4)^39^ was run to call structural variants (SVs), using default settings. Only SVs larger than 1kb were considered. SVs 1-30 kb in size needed quality scores ≥300 but those larger than 30 kb used a cutoff of 250. SVs required breakpoint assembly on both sides with at least four discordant and two split reads supporting them. Where breakpoints were imprecise, defined as the start and end positions being >10bp apart, the SV was filtered out. SVs where the standard deviation of the alignment position at the ends of discordant read pairs was <5 were also removed. Lastly, to remove potential germline SVs and recurrent artefacts, we removed SVs found in at least three different samples from a panel of normal samples or in the matched normal sample^40^.

### Variant calling and filtering for duplex sequencing of the *NF1* gene

For each patient, a matched normal was created by aggregating samples from the same patient. Patients PD50297, PD51122, and PD51123 had sufficient samples contributing to their matched normals that germline mutations could be confidently removed and no hard VAF cutoff was applied. Samples from the validation cohort had fewer tissues per donor, and so any mutation at an allele fraction of greater than 0.1 in the deduplicated bam was excluded on the grounds that it may be a germline variant.

Reads were collapsed into read bundles that originate from the same duplex DNA molecule using their adaptor sequences. Mutations were called if they were supported by reads from both strands and were not called in the normal sample^21^.

Duplex sequencing is vulnerable to trace amounts of contamination as a result of its ability to detect mutations on a single molecule of DNA. Contaminating DNA may be either human or non-human. In order to filter out non-human contamination when calling substitutions, only read bundles that had either 0 or 1 single-stranded or double-stranded mismatches relative to the reference genome in addition to the called substitution were included in the analysis. This filters out contamination with DNA from other species, which may have multiple mismatches per read bundle, as well as reads with large amounts of single-stranded DNA damage. Contamination with human DNA may be either from samples that are on the same plate (either biological contamination through errant DNA molecules or bioinformatic contamination through index hopping), or from DNA that is not on the plate. To remove contaminants from the same plate, every sample on the plate was genotyped both with GATK HaplotypeCaller^41^ and BCFTools^42^, and no mutation that was called as germline in any sample was considered for a given plate. Second, any mutation catalogued as germline in Gnomad v2^43^ with an allele fraction of greater than 0.1% was removed from the analysis. After these steps, the global dNdS value for the dataset was 0.93 (95% CI 0.85-1.03). Reassuringly, contamination with other species or germline DNA from other humans contains an excess of synonymous mutations and artificially lowers the dN/dS ratio. Any contamination that passes our filters will reduce our ability to detect positive selection, but is highly unlikely to contribute to detection of positive selection where there is none.

After filtering, mutations were analysed using the package dNdScv^15^ . This tool uses a maximum likelihood method to quantify selection, integrating genomic covariates to correct for mutation biases. No limit was applied to the number of mutations with the *NF1* gene or the number of mutations per sample, and the duplex sequencing coverage of *NF1* in the group being tested was used as an offset.

One variant in *NF1*, chr 17:31226459 A>AC resulting in p.I679fs, was called by duplex sequencing in five separate tissues from PD51122 at low allele fraction: the pons (VAF 0.02214), the medulla (VAF 0.00744), the occipital cortex (VAF 0.00096), the small bowel (VAF 0.00147), and a spindle cell lesion of the ankle (VAF 0.00059). It was also previously found by whole genome sequencing of the pons and genotyped in the medulla. Four possible explanations for this recurrence presented themselves:

i. Artefact;
ii. Carryover. It occurred in only one tissue, and other tissues were contaminated with DNA from this tissue;
iii. Embryonic. The variant occurred in a common ancestor of cells in all of these tissues; or
iv. Independent. It occurred independently in each tissue.

To address each in turn: (i) The variant was called by whole genome sequencing and duplex sequencing. The latter has an extremely low error rate. The variant has only been found in the patient with neurofibromatosis type 1, and so is unlikely to represent a mapping error or similar. All occurrences of this variant are also out of phase with the germline *NF1* variant. It is therefore unlikely to be an artefact. (ii) If material from one tissue was contaminating another, numerous mutations from the source tissue should be found in the contaminated tissue rather than just the *NF1* variant. Additionally, we would not expect to find carryover mutations in laser capture microdissected biopsies, since this tissue has been directly visualised and stray cells would be apparent. Assuming that the pons is the source of the mutation (as the mutation was found here by whole genome sequencing, and the VAF is highest here), we can look for pontine variants in other tissues. Looking in the medulla, for instance, of 110 pontine substitutions, 14 were found in the medulla. Of these, 11 were present in laser capture microdissections of the medulla. Only 3 mutations, therefore, are consistent with carryover. This is an implausibly low number to suggest carryover. (iii) It is highly unlikely that a genuine embryonic variant that has occurred across developmentally distantly related organs such as the bowel and the central nervous system would be at such low allele fraction in these different tissues. Such a finding would imply a cell that existed late in development that could give rise to both bowel and brain. For the mutation that we identified across more than one germ layer in whole genome sequencing data (Y489C), our analysis of other shared mutations (**Fig. 2e**) indicated that they were independent events and not embryonic. Furthermore, the spindle cell lesion of the ankle is clonal. An embryonic variant within the neoplasm, therefore, should be clonal within it, and not at the low VAF that we observe. (iv) On first inspection, the same precise DNA changes are unlikely to occur multiple times. The mutation, however, involves slippage in a cytosine homopolymer, which may be highly mutable. Exactly the same mutation has occurred in 60 different cancers as documented in Cosmic^45^, such that this locus has far more indels than any other in the *NF1*. The high dN/dS values associated with *NF1* truncating mutations on a background of neurofibromatosis type 1 suggest that essentially every time a truncating variant occurs in *NF1* it is likely to be selected for, and so we have a good chance of finding it.

A combination of the above explanations is also possible. For example, the mutation may be a shared embryonic variant between the pons and medulla, but has occurred again independently in the ankle lesion. Following the reasoning above, we considered independent mutations to be most likely and the results presented in the main text treat them as such. The dN/dS analysis quantifying selection pressures in *NF1* has been repeated both treating the mutations as independent and only allowing each variant to be counted only a single time in each patient. The conclusions drawn from the data remain the same using both methods (**Supplementary figure 9**).

### Tumour phylogeny reconstruction

Phylogenies were created from the bulk tumour samples with purity ≥40% only, to ensure confidence in the copy number calling estimates. Battenberg copy number calls were used to generate cancer cell fractions (CCFs) for the substitutions which were then subject to multidimensional DPClust (ndDPClust, implemented using DPClust version 2.2.2)^32,44^. The Gibbs sampler was run for 1,000 iterations with 200 dropped as burn-in per tumour. The resultant mutation clusters were then inspected and kept, split or merged depending on the distribution of the CCFs of their substitutions across all samples. Only clusters containing at least 1% of substitutions were kept (**Supplementary table 10**). The final clusters demonstrated the clonal trunk and multiple subclones found across each tumour.

### Estimating tumour infiltration

The truncal mutations (both driver and passenger mutations) identified during tree building provided a set of variants that would be expected in any sample infiltrated by tumour. All samples from each patient, irrespective of their distance from the tumour, were examined for evidence of tumour infiltration. For the two near-diploid tumours (PD50297 and PD51123), we identified any truncal variant found at a site with one clonal copy of each allele in all bulk tumour samples with a purity ≥40%, resulting in 384 and 259 clonal substitutions respectively. The mean VAF, excluding outliers, of these variants was calculated and doubled to provide an estimate for tumour infiltration. Even a normal sample that contains no tumour cells may share early embryonic variants with a tumour, and it is important to distinguish embryonic variants from truncal tumour variants; this may be done based on the overall distribution of the VAFs in the sample. A normal sample with no tumour infiltration will share a handful of high VAF variants with the tumour (representing embryonic variants shared by the tumour and normal tissue) whilst a sample with low level tumour involvement will possess many truncal variants at low VAFs. To remove outlier embryonic variants, VAF vectors were trimmed to only include variants whose VAF in a given sample was no further than 1.5× interquartile range below the lower or above the upper quartile. Any variant outside of this range was deemed an outlier. For the grossly aneuploid tumour (PD51122), the substitutions at uniformly clonal 2+0 copy number sites that were estimated to be found on both copies of the allele were used to estimate tumour infiltration; no doubling of the VAF was required. PD51122 had 52 substitutions that fulfilled these criteria.

This approach was evaluated using two methods. First, for the bulk tumour samples with a purity ≥10%, the minimum threshold typically required for copy number callers, we could compare our purity estimates with those generated by Battenberg. There was a near perfect, positive correlation in the results between the two methods (**Supplementary figure 7B**). Second, to establish the efficacy of our method for samples with <10% tumour involvement, we simulated data to capture a wide range of trunk lengths, sample coverage and tumour involvement. For each simulation (1,000 in total per scenario), the total coverage for each locus in the tumour trunk was randomly sampled from a Poisson distribution with a mean equal to the average sample coverage. The variant depth was generated from a binomial distribution whose number of trials was equal to the coverage generated by the aforementioned Poisson distribution and probability of success was half the purity, like the scenario used for the near-diploid tumours. The fraction of simulations, for a given number of truncal mutations, coverage and tumour purity, that returned a tumour infiltrate value >0, provided a probability for detecting tumour, if there was genuine tumour involvement in a sample. In the worst case scenario, with only 50 substitutions to use in the pile-up, we estimated we had >95% chance to detect 3% tumour infiltration and 100% chance at 5% infiltration using our approach for a sample with 30× coverage (**Supplementary figure 8**).

For the whole exomes, where the baitset would not capture most truncal mutations identified in the whole genomes, the tumour purity was approximated using the variant allele fraction of each tumour’s *TP53* hotspot mutation, correcting for its copy number state.

### Driver mutation identification and annotation - whole genome and whole exome sequencing - substitutions and indels

For substitutions and indels, all mutations resulting in protein coding changes in genes reported in the COSMIC (version 94) cancer gene census were initially considered^45^.

Driver mutation status was assessed prior to the application of the exact binomial filter (which determines germline status - see above). This circumvented the risk that true driver mutations might be eliminated at a subsequent filtering step, e.g. germline driver events. Two classes of mutations were considered to be candidate drivers: first, missense substitutions or in-frame indels occurring at hotspots in dominant-acting genes; and second, mutations in recessive-acting cancer genes predicted to result in loss of function, such as nonsense, frameshift, or essential splice site variants. Candidate mutations were assigned to tiers, according to their likelihood of acting as a driver. Tier 1 substitution/indel drivers occurred within genes that were recurrently mutated in a recent meta-analysis of over 1,000 paediatric high-grade gliomas^16^. This list of genes included: *ACVR1*, *ASXL1*, *ATM*, *ATRX*, *BCOR*, *BRAF*, *CCND2*, *CDK4*, *CDK6*, *CDKN2A*, *CDKN2B*, *EGFR*, *FGFR1*, *H3F3A*, *HIST1H3B*, *HIST1H3C*, *HIST2H3C*, *ID2*, *KDM6B*, *KDR*, *KIT*, *KRAS*, *MET*, *MYC*, *MYCN*, *NF1*, *NTRK1*, *NTRK2*, *NTRK3*, *PDGFRA*, *PIK3CA*, *PIK3R1*, *PPM1D*, *PTEN*, *RB1*, *SETD2*, *TERT*, *TOP3A*, *TP53*. Tier 2 mutations occurred in other supposed cancer genes from the COSMIC (version 94) cancer gene census list.

Mutations in *NF1* itself were considered differently. Inactivating mutations were considered to be probable drivers. Although NF1 is a recessive cancer gene, it does have residues that are mutated more frequently. Missense mutations that occurred in such loci, defined as >4 mutations of a given residue in COSMIC, were considered as probable driver mutations, and further support for their functional effect was sought from the literature and from predictors of mutational effect^46^.

### Driver mutation identification and annotation - whole genome and whole exome sequencing - copy number changes

Copy number changes were determined to be driver events according to sample ploidy, the genes found on each segment and the segment length. Oncogenes were considered to be amplified if their total copy number was ≥5 when ploidy was <2.7 or ≥9 when ploidy was ≥2.7. For tumour suppressor genes, total copy number had to equal 0 for <2.7 ploidy and ≤ (ploidy - 2.7) when ploidy ≥2.7. Copy number aberrations passing these criteria were then annotated as putative tier 1 driver mutations if the oncogene(s) or tumour suppressor gene(s) they contained was found in the list of genes above, the segment width was ≤10Mb wide and this was a recognised oncogenic event for that gene. For example, copy number changes in genes that mediate oncogenesis via fusion events alone were not considered Tier 1 drivers. Tier 2 drivers did not meet the criteria outlined for Tier 1 variants, but had to be found on segments ≤1Mb wide.

### Driver mutation identification and annotation - whole genome and whole exome sequencing - structural variants

For a structural variant, independent of copy number state, to be annotated as a driver, it had to either form a fusion gene recognised to be oncogenic, truncate the gene footprint of a tumour suppressor gene or activate an oncogene through intragenic deletion (e.g. *PDGFRA*). Once again, Tier 1 events occurred in the list of genes used for other variant classes whereas Tier 2 events were plausible drivers that fell outside of these.

### Testing for recent clonal expansions associated with *NF1-*null status

A linear mixed effects model comparing the mutation burden derived from whole genome sequencing of *NF1*-null vs *NF1*-heterozygous histologically-normal central nervous system biopsies/microbiopsies was fitted in R using the package nlme. *NF1*-null status, whether the sample was derived from bulk sequencing or laser capture microdissection, and coverage were included as fixed effects. The piece of tissue from which the sample was derived was used as a random effect (i.e. two microbiopsies from the same piece of tissue should be correlated with one another). Although *NF1*-null status was associated with a statistically significant effect, the effect size was only of seven additional mutations. Given that the postnatal somatic mutation rate of most tissues is 10-50 mutations per year (including a rate in glia of 27 substitutions per year^47^ and a rate in neurones of 17 substitutions per year^21,47^, and the prenatal rate is usually higher, a recent clonal expansion should result in a mutation burden on the order of 100 mutations even in a child; we therefore concluded that *NF1-*null status was incompatible with a recent clonal expansion.

### Detecting independent *NF1* null clones in normal tissue - whole genome and whole exome sequencing

Driver events within *NF1* were initially identified in the same manner as all other driver mutations. This included a germline essential splice site *NF1* mutation within PD51122 that accounted for their Neurofibromatosis type 1. Second loss of function *NF1* mutations in this case were assumed to render the affected cells “*NF1*-null”.

Evidence for any *NF1* driver point mutation that had been identified in the child with neurofibromatosis type 1 was sought in the remaining two cases in the discovery cohort. Similar to during the substitution filtering, this provided an approximation of the base sequencing error rate, above which we could determine an *NF1* point mutation to be truly present in the index case. We performed a one-sided Fisher’s exact test using the summed variant and total read depth from the two children without *NF1* mutation against those observed in each microdissection and bulk biopsy from the child with neurofibromatosis type 1. After a multiple hypothesis testing correction (Benjamini-Hochberg method), the *NF1* point mutation was considered present in a sample if *q* <0.01. All *NF1* driver point mutations that were identified were found in at least one sample without any detectable tumour involvement. No copy number aberrations or structural variants involving *NF1* were detected in normal tissues using standard variant calling.

To increase our sensitivity to detect loss of heterozygosity (LOH) events in the normal tissue of the child with neurofibromatosis type 1, we phased SNPs to each gene allele. The child’s tumour had LOH of the entirety of chromosome 17, leaving only copies of the allele bearing the germline *NF1* mutation. We could phase the heterozygous SNPs identified in the deeply sequenced blood sample (PD51122q) on chromosome 17 according to which allele had the greatest allele frequency in one of the purest bulk tumour samples (PD51122m). These phased SNPs were then profiled in all remaining samples. Only SNPs with ≥10× coverage in a sample were kept for its downstream analysis as few SNPs in non-coding regions would be captured by whole exome sequencing.

To identify samples with possible independent LOH events inactivating *NF1* in both the whole genome and exome sequencing data, a two-sided exact binomial test was performed. In this test, the number of trials was the sum of total coverage across the heterozygous SNPs found across the gene. The number of successes was the sum of the depth of the alleles that were only found in the tumour. The hypothesised probability of success was the expected aggregate allele fraction. The *NF1* locus was 2+0 in the tumour, while a normal cell would have one copy of each parental allele. The aggregate allele fraction therefore would equal *tumour purity + ((1 - tumour purity) x 0.5)*. *P* values underwent multiple hypothesis correction using the Benjamini-Hochberg method. To ensure confidence that we were truly detecting these in normal tissue, only samples where *q* < 0.01, the median coverage was ≥30× and tumour purity was <1% were considered to possess a copy number change to the *NF1* locus that could not be explained by tumour infiltration alone.

Two samples that had significant shifts in the proportion of each *NF1* allele unusually favoured the wildtype allele (PD51122s_lo0012 and PD51122u_lo0009). These microdissections of non-neuroectodermal origin are interpreted as containing clones with LOH events that lost the mutant *NF1* allele.

The *NF1* locus had not undergone LOH in the other two children, meaning that a similar analysis could not be performed.

### Assessment of the genetic relationship between tissues that shared a somatic *NF1* variant in PD51122

For a variant to be found in two tissues, either it must have been acquired from a shared ancestor or developed independently. Assuming a comparable rate of mutation acquisition, tissues with a more recent common ancestor will share a greater number of mutations than those that are more distantly related. To determine whether the tissues carrying the same somatic *NF1* mutation were uniquely related, we first needed a control to determine how related two tissues would be by chance in this child.

To construct our control data, we used the normal tissues (<1% estimated tumour contamination) without evidence of a second *NF1* hit. Separate comparisons of normal CNS vs normal CNS and normal CNS vs normal mesoderm were made to account for differences in the genetic architecture between germ layers and tissues. A mutation was determined to be shared between tissues if it was identified in both using the aforementioned Shearwater-like approach, rather than relying on the calls from the variant caller alone. This improved our sensitivity for detecting low VAF variants and mitigated some of the risk that true shared variants would not be called or erroneously filtered in one sample by our pipeline. All samples in each group were then iteratively compared to the others.

The pairwise comparison was then repeated for tissues that shared a somatic *NF1* variant and the mean number of shared substitutions per pair was calculated for each mutation (test data). The same number of pairwise comparisons for each mutation were then drawn from the control data at random, without replacement, and the mean calculated. This was repeated 1,000 times. The *P* value was determined by the number of draws where the control data mean was greater than that observed in the test data (one-sided test).

## DATA AVAILABILITY

Whole-genome and targeted sequencing data are being deposited in the European Genome-Phenome Archive (EGA; https://www.ebi.ac.uk/ega/) with accession ID EGAD00001015398. Mutation calls are available as supplementary tables.

## CODE AVAILABILITY

Custom R scripts used to analyse the data are available from the authors upon request.

## ACKNOWLEDGEMENTS

This study was funded by the Wellcome Trust (institutional grant; personal fellowships to T.R.W.O. and S.B.; grant number 206194, 108413/A/15/D & 223135/Z/21/Z). Additional funding was received from: The Brain Tumour Charity (T.S.J.) directly and via the INSTINCT network and EVEREST centre; Children with Cancer UK; Great Ormond Street Hospital Children’s Charity; Olivia Hodson Cancer Fund; Cancer Research UK; National Institute of Health Research; NIHR Biomedical Research Centres (Great Ormond Street Hospital and Cambridge University Hospitals); The Royal National Orthopaedic Research and Development Department (A.M.F.); The Bone Cancer Research Trust (A.M.F.). We thank the CCLG Tissue Bank, the CCLG centres and the ECMC Paediatric Network for the collection and provision of tissue samples (project number 2016 BS 05). The CCLG Tissue Bank is funded by Cancer Research UK and CCLG. A.M.F. is also separately supported by the National Institute for Health Research, UCLH Biomedical Research Centre and the UCL Experimental Cancer Centre. Funding from these institutions supported the work of the Biobank where the samples from the adult cohort were stored. We thank the clinical teams of Cambridge University Hospitals and Great Ormond Street Hospital, including the mortuary staff. Vanna Lee, Lakiesha Ward and Oluminde Ogunbiyi from Great Ormond Street Hospital and Aimee Whyte from Addenbrooke’s Hospital helped facilitate the collection and transfer of samples for which we are grateful. We thank Giulio Caravagna for his assistance with copy number calling quality control and Aidan Maartens (science writer at the Wellcome Sanger Institute) for his critical review of the manuscript. We are indebted to the families who participated in this research.

## AUTHOR CONTRIBUTIONS STATEMENT

S.B. and T.R.W.O. designed the experiment and co-wrote the manuscript. T.S.J. and K.Al. conducted the autopsies and provided neuropathological histology expertise. T.R.W.O. performed microdissection, with laboratory support provided by Y.H. and P.N., with further technical support from A.T. T.R.W.O., A.R.J.L., and H.L-S. analysed data, with the assistance of R.S., M.D.Y., T.H.H.C., H.J.,

T.M.B. and I.C. M.D.Y. provided statistical expertise. A.B., K.Aq. D.H., M.J. and T.D.T. provided clinical expertise and contributed to discussions. U.L. provided radiological expertise. F.A.J. and A.M.F. provided pathological expertise. S.B., T.S.J., I.M. and K.Al. co-directed the study.

## EXTENDED DATA

**Supplementary Fig. 1:**
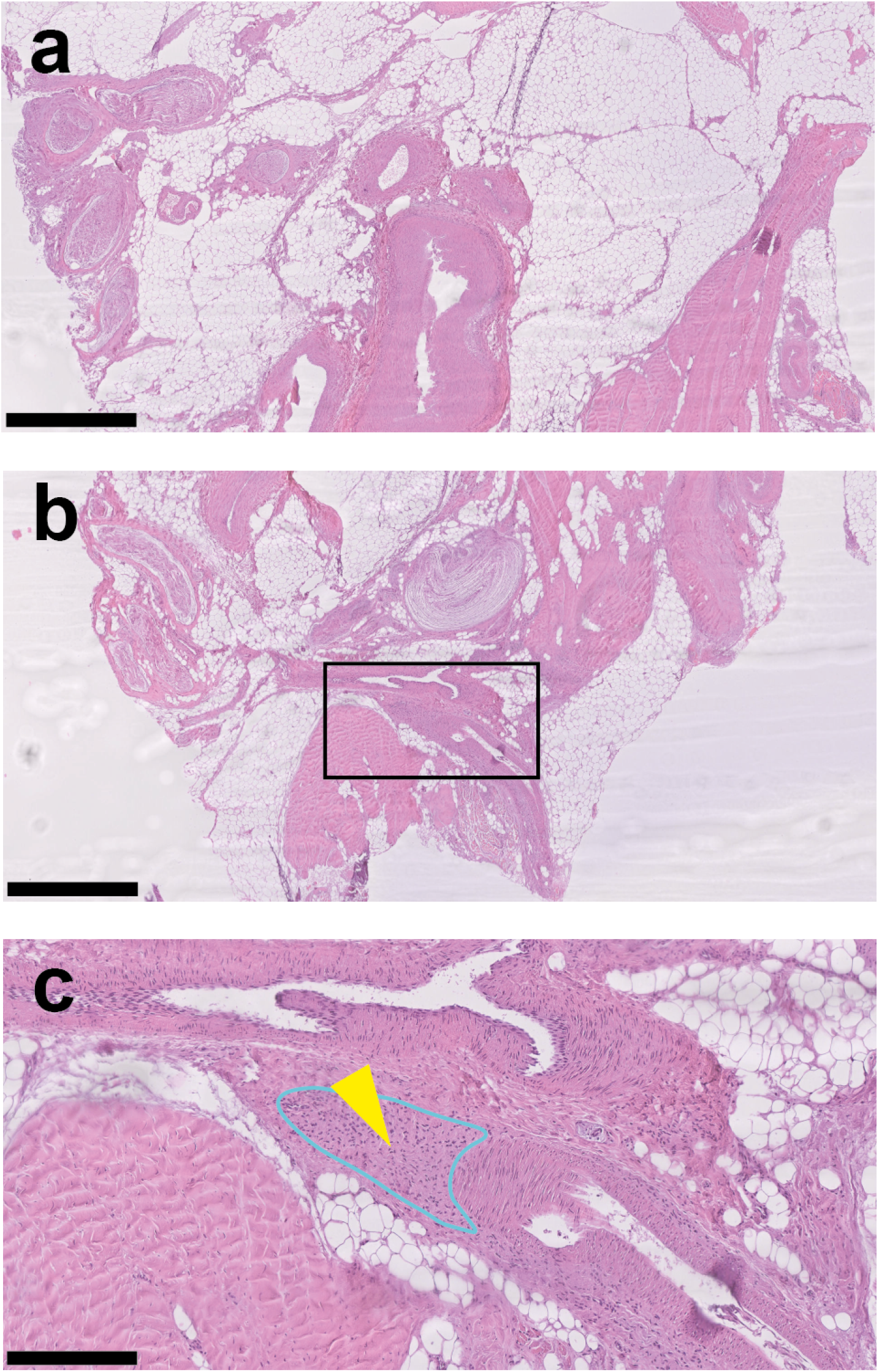
Subcutaneous ankle lesion in the child with neurofibromatosis type 1. Representative histological appearances of the tissue are shown (**a**, **b**). The tissue comprises adipose tissue, bands of fibrous tissue, a Pacinian corpuscle, thick nerve trunks and ganglia. The light blue outline with a yellow arrowhead in **c** indicates the region microdissected that yielded a biallelic loss of *NF1* (PD51122ac_lo0013). It comprises bland spindled cells, embedded in fibrous and fibrillary stroma, that surround a blood vessel. The scale bars represent 1mm, 1mm and 250 µm respectively.

**Supplementary Fig. 2:**
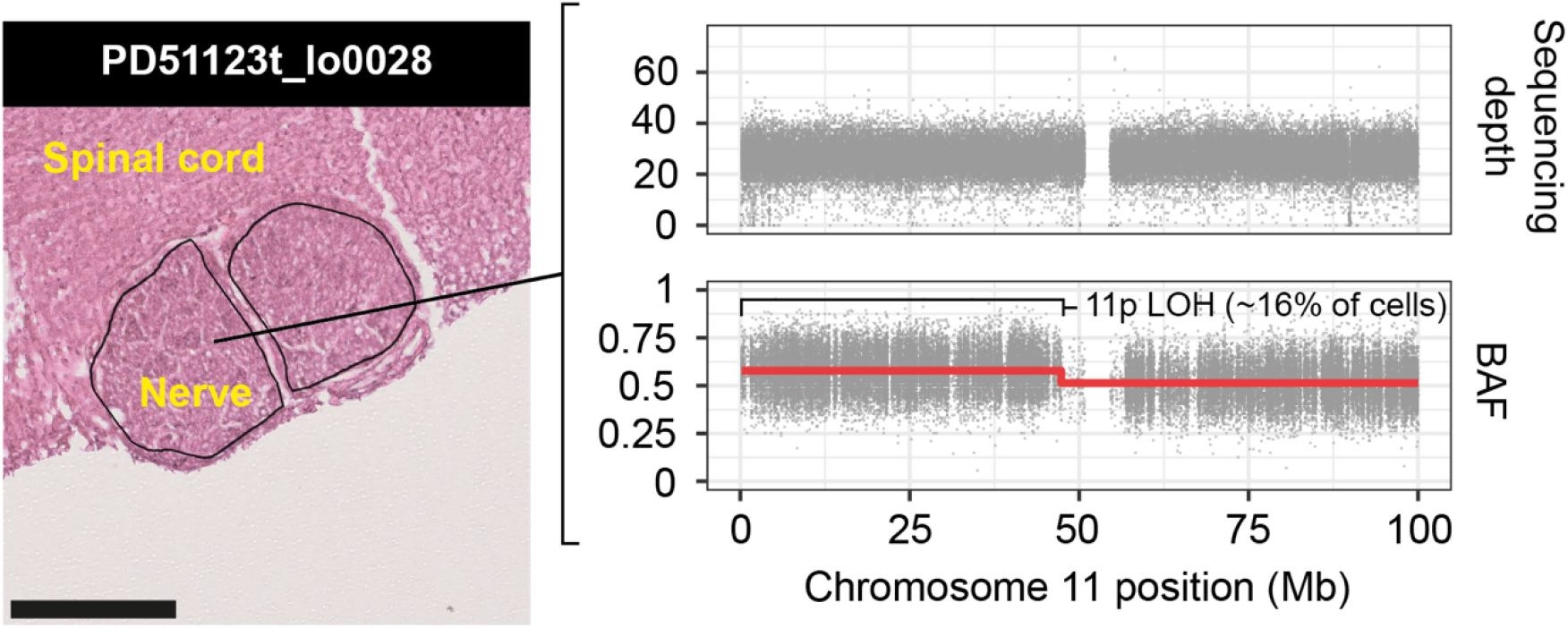
Mosaic uniparental disomy of chromosome 11p in a nerve from case PD51123. Left, a histological image of the slide from which the affected sample was microdissected. Scale bar represents 250 µm. Right, the coverage of chromosome 11 SNPs and their B-allele fraction (BAF). The B-allele here is the one inferred to be derived from the major allele within predicted haplotype blocks (red line). The BAF split was not detected by ascatPCA but detected on manual review and the output here is from Battenberg.

**Supplementary Fig. 3:**
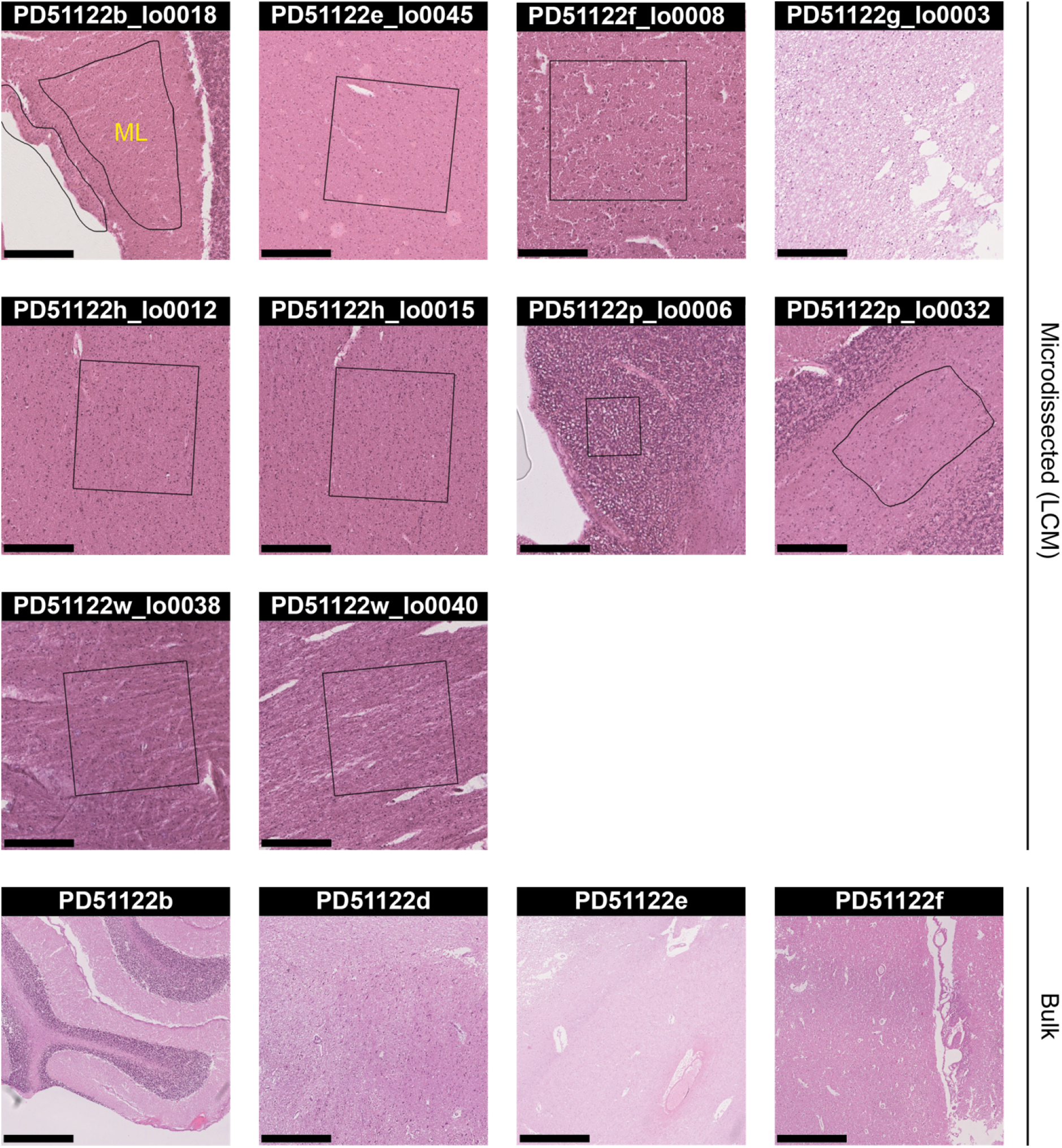
Histological images of normal tissues with independent *NF1* null clones. Where the clone was detected in a microdissected sample, the exact section and area sequenced is shown. For clones found in bulk samples, a representative image from the reference slide is provided. The microdissected and reference sections are 16 µm and 4 µm thick respectively. PD51122g_lo003 was taken from a section which had dried prior to slide scanning and microdissection, resulting in a suboptimal image. Consequently, the reference slide was used here to provide an overview of the approximate area microdissected. The black outlines on images represent the perimeter of microdissection. PD51122b_lo0018 is a region of the molecular layer (ML), taken from the cerebellum. The scale bars for the images of microdissected samples represent 250 µm. The scale bars for PD51122b, PD51122e and PD51122f are 1 mm and for PD51122d 500 µm.

**Supplementary Fig. 4:**
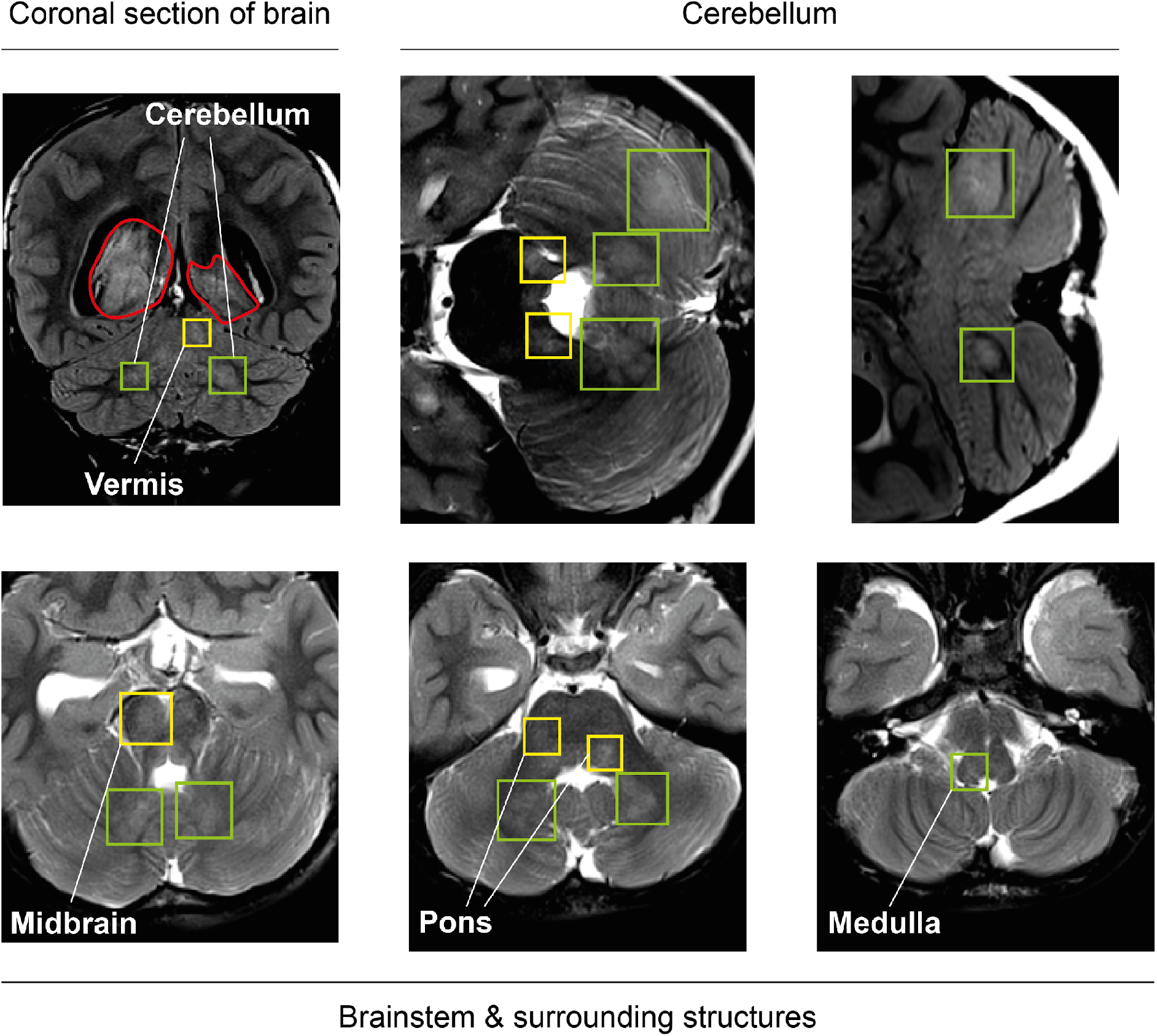
Brain MRI at diagnosis from the child with neurofibromatosis type 1. Coronial FLAIR (top row, left and right images) and axial T2-weighted images (all other images; top row, middle and right images are rotated 90° anticlockwise) show typical focal areas of signal intensity (FASI) in the white matter and cerebellar cortex (green boxes). The presence of tumour (red outline) makes it challenging in some areas to distinguish tumour from FASI (yellow boxes). No convincing FASI were found in the cerebral cortex or spinal cord, suggesting that no close correlation of the *NF1* null clones and FASI could be made. Note that there is hyperintensity of both hippocampi (bottom row, left image).

**Supplementary Fig. 5:**
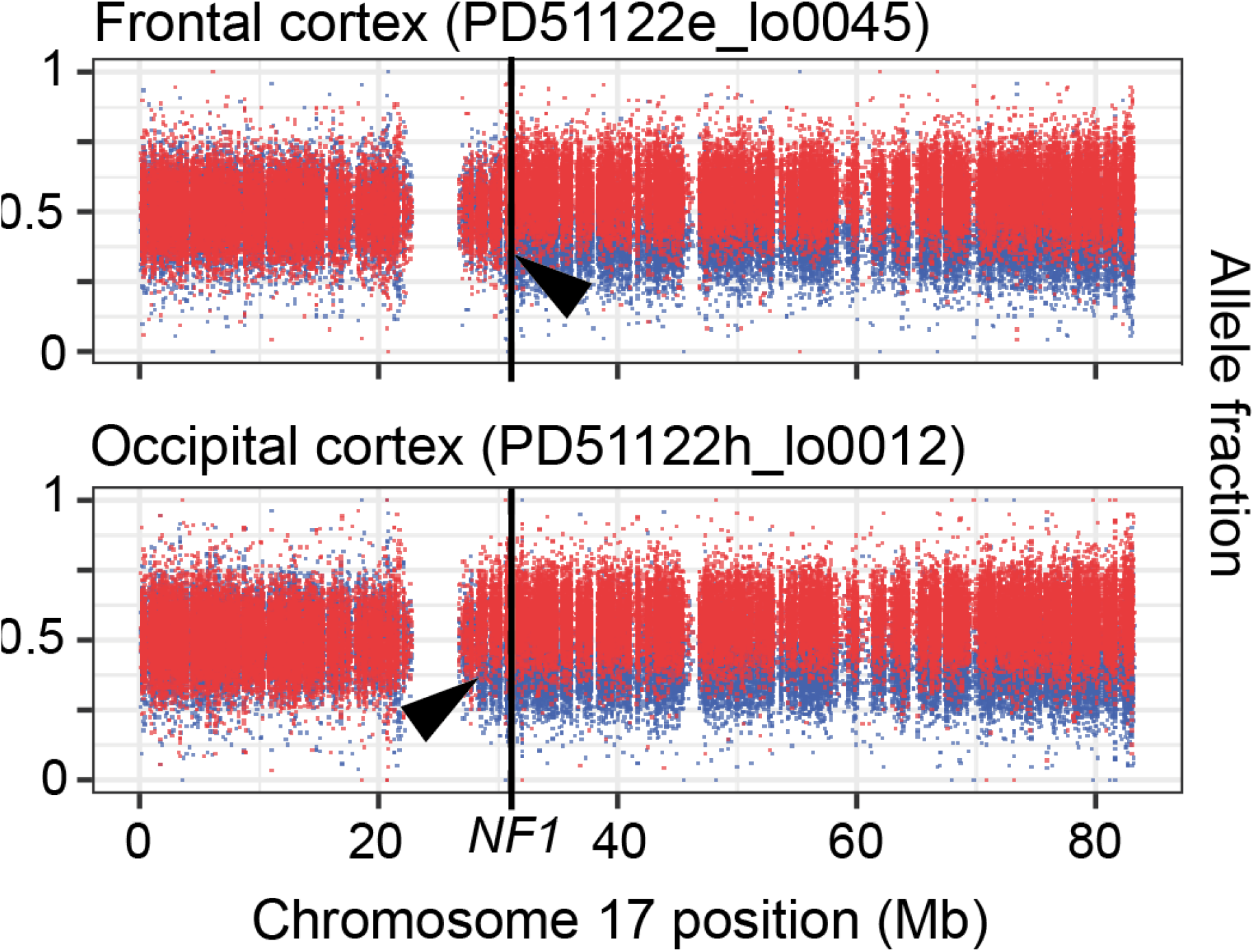
Allele fraction plots for the normal tissues in the child with neurofibromatosis type 1 with the largest *NF1* LOH events. Each dot represents a heterozygous SNP that is phased to either the copy of chromosome 17 bearing the germline NF1 mutation (red) or the wildtype allele (blue). Black arrowheads indicate the approximate location of the breakpoint in each sample.

**Supplementary Fig. 6:**
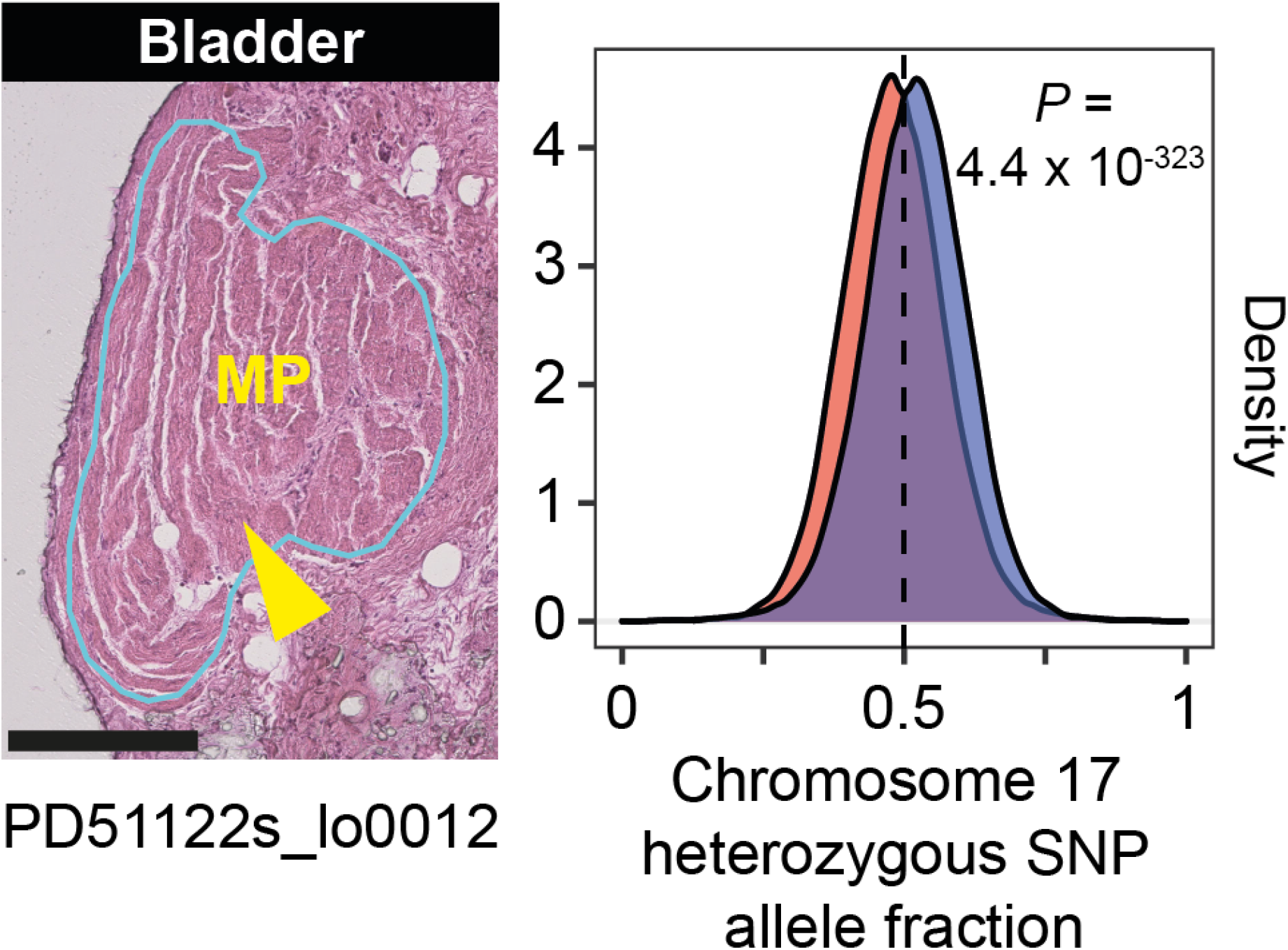
Loss of the germline mutated NF1 allele in bladder muscle. Left, a histological image of the tissue microdissected to generate PD51122s_lo0012. Right, a density plot for the two alleles at heterozygous SNP sites found across chromosome 17. A two-sided, exact binomial test is applied between the observed chromosome 17 allele fraction bearing the germline mutant *NF1* (red) and the expected fraction (dashed line) (**Methods**). The scale bar represents 250µm. MP, muscularis propria.

**Supplementary Fig. 7:**
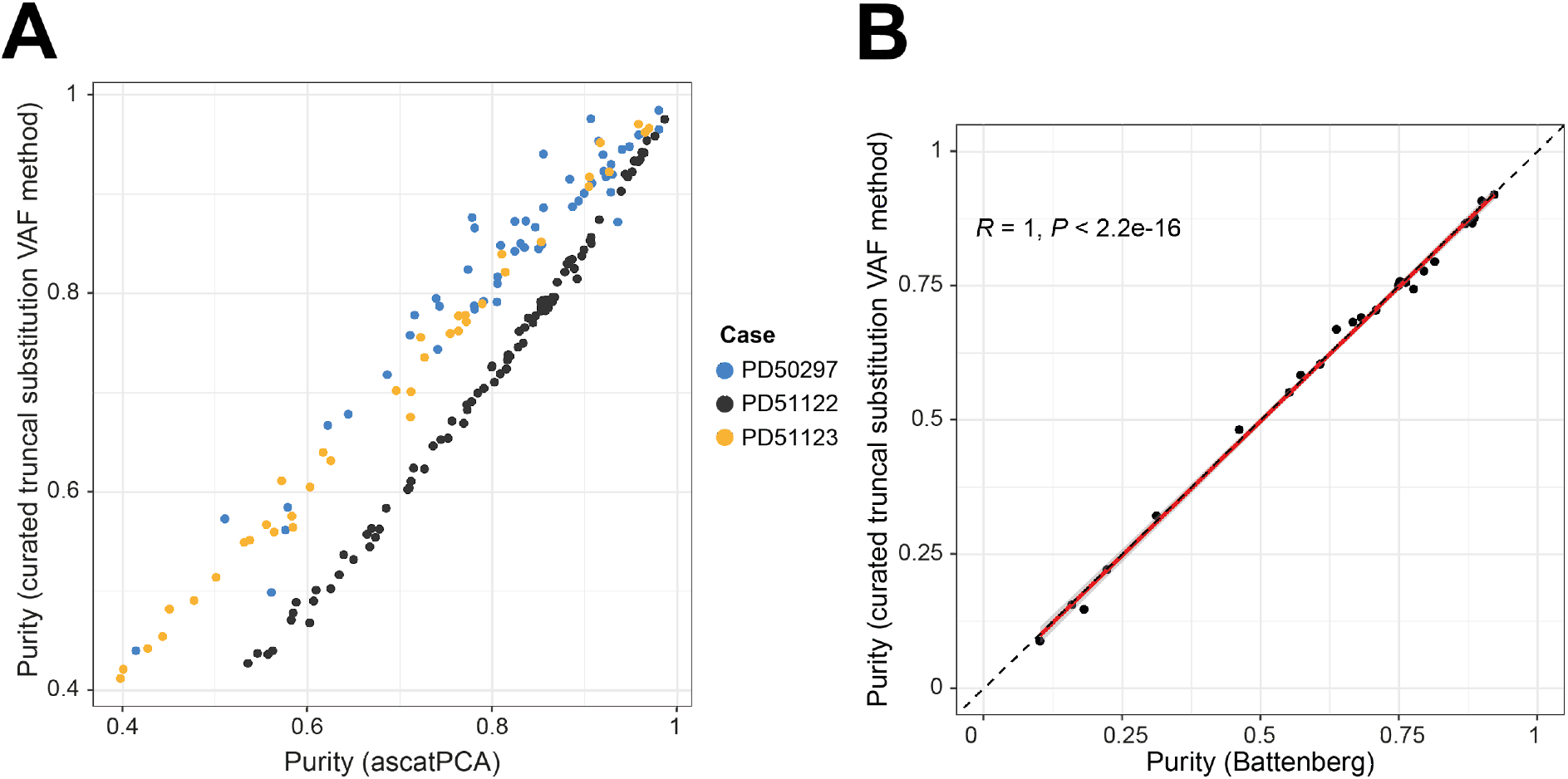
Comparison of tumour purity estimates between three methods. **(A)** Our truncal mutation VAF method versus ascatPCA, restricted to tumour samples with a tumour purity ≥40% (as determined by the former method). **(B)** Our method versus Battenberg, restricted to bulk tumour samples with a tumour purity ≥10% (as determined by Battenberg). A Pearson correlation between the two axes is shown on the graph. A linear regression line is fit to the data (red, solid), with 95% confidence intervals (grey ribbon). It closely matches a line representing the equation y = x (black, dashed).

**Supplementary Fig. 8:**
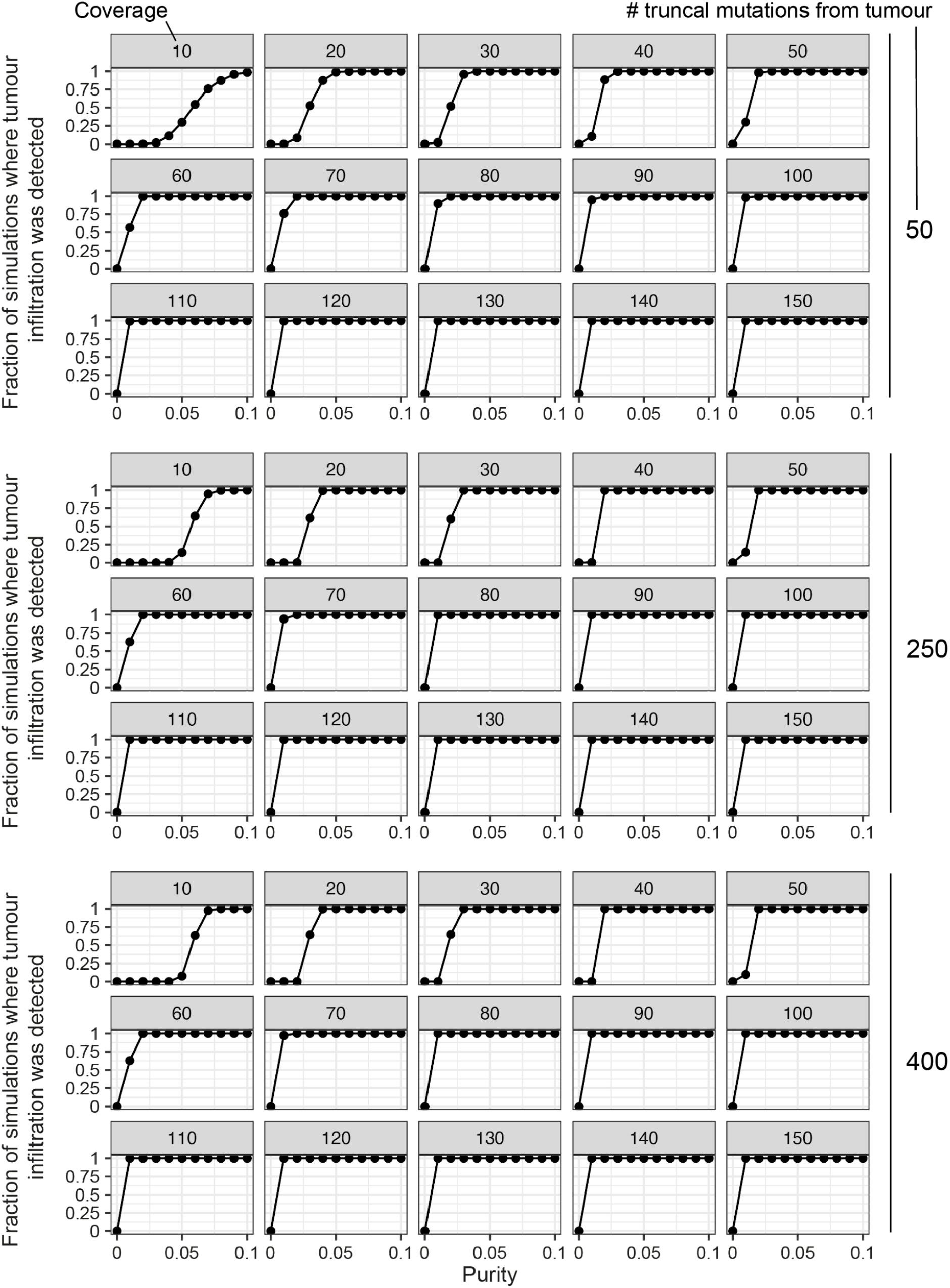
Simulated data estimating the efficacy of our approach to detect low level tumour infiltration. The range of truncal mutation catalogue sizes used reflects the variability seen in our cohort.

**Supplementary Fig. 9:**
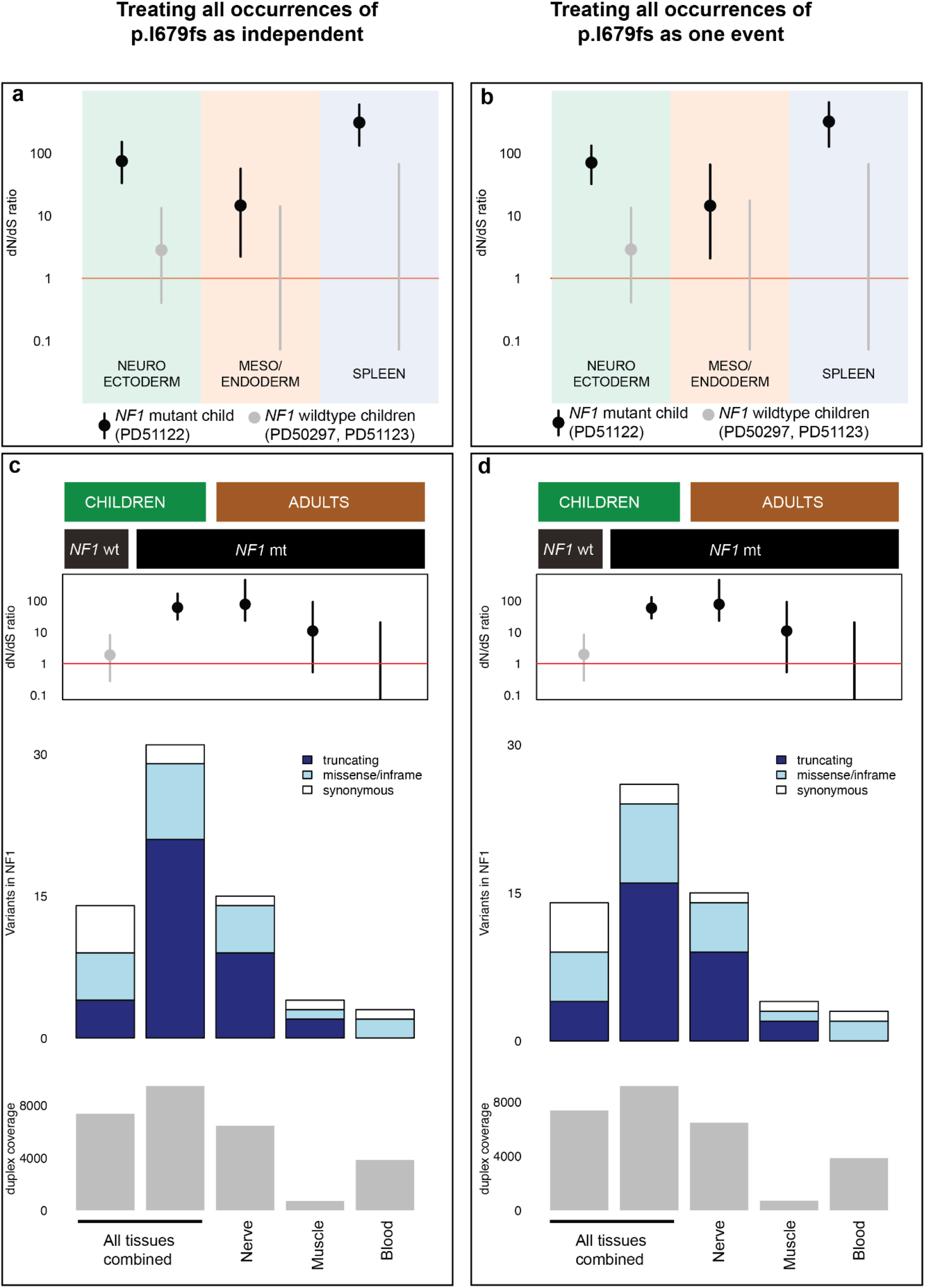
Duplex sequencing data analysed considering the same mutation as independent or as the same event. One variant in *NF1*, chr 17:31226459 A>AC resulting in p.I679fs, was called by duplex sequencing in five separate tissues from PD51122. These may have occurred as independent mutations or may represent the same event (Methods). Here, the results are shown comparing each occurrence of this mutation as independent events (panels **a** and **c**; as in Fig. 2f and g) or as the same event (panels **b** and **d**). **a** and **b**, dN/dS ratios for truncating variants, according to germ layer and *NF1* germline mutation status. The dot represents the maximum likelihood estimate and the black lines represent the 95% credible interval. When the lower bound of the credible interval is above 1 (red line), there is statistically significant positive selection. **c** and **d**, normal tissues from adults with neurofibromatosis type 1 are grouped by tissue type and evaluated for an excess of non-synonymous variants in *NF1*, and compared with the three index children. Upper, dN/dS ratios for truncating mutations; middle, counts of variants in *NF1*; lower, total duplex coverage (**Methods**) over *NF1* in each group.

